# Regulation of the mouse ventral tegmental area by melanin-concentrating hormone

**DOI:** 10.1101/2023.09.25.559435

**Authors:** C Duncan Spencer, Persephone A Miller, Jesukhogie G Williams-Ikhenoba, Ralitsa G Nikolova, Melissa J Chee

## Abstract

Melanin-concentrating hormone (MCH) acts via its sole receptor MCHR1 in rodents and is an important regulator of homeostatic behaviors like feeding, sleep, and mood to impact overall energy balance. The loss of MCH signaling by MCH or MCHR1 deletion produces hyperactive mice with increased energy expenditure, and these effects are consistently associated with a hyperdopaminergic state. We recently showed that MCH suppresses dopamine release in the nucleus accumbens, which principally receives dopaminergic projections from the ventral tegmental area (VTA), but the mechanisms underlying MCH-regulated dopamine release are not clearly defined. MCHR1 expression is widespread and includes dopaminergic VTA cells. However, as the VTA is a neurochemically diverse structure, we assessed *Mchr1* gene expression at glutamatergic, GABAergic, and dopaminergic VTA cells and determined if MCH inhibited the activity of VTA cells and/or their local microcircuit. *Mchr1* expression was robust in major VTA cell types, including most dopaminergic (78%) or glutamatergic cells (52%) and some GABAergic cells (38%). Interestingly, MCH directly inhibited dopaminergic and GABAergic cells but did not regulate the activity of glutamatergic cells. Rather, MCH produced a delayed increase in excitatory input to dopamine cells and a corresponding decrease in GABAergic input to glutamatergic VTA cells. Our findings suggested that MCH may acutely suppress dopamine release while disinhibiting local glutamatergic signaling to restore dopamine levels. This indicated that the VTA is a target of MCH action, which may provide bidirectional regulation of energy balance.

**Significance Statement:** Role of melanin-concentrating hormone (MCH) on energy balance may converge on the dopamine system via the mesolimbic pathway, as loss of MCH or MCH receptor (MCHR1) signaling increases hyperactivity and energy expenditure associated with a hyperdopaminergic state. MCH can suppress dopamine release within the mesolimbic pathway, but its underlying mechanism is not known. We thus determined if MCH could inhibit dopamine release through direct actions within the ventral tegmental area (VTA). We found that MCH directly inhibited dopaminergic VTA cells, but MCH also disinhibited excitatory input to dopamine cells. Therefore, we showed that the VTA is a putative target site supporting dopamine-dependent actions of MCH.

## Introduction

Melanin-concentrating hormone (MCH) is produced largely in the lateral hypothalamic zone to promote positive energy balance by acting through its receptor MCHR1. MCHR1 is a G-protein coupled receptor (Chambers et al., 1999; Lembo et al., 1999; Saito et al., 1999) and typically inhibits cells via G_i/o_ activation (Gao & Pol, 2001). Mice overexpressing *Mch* may overeat and develop diet-induced obesity (Ludwig et al., 2001), while *Mch* knockout mice (*Mch-KO*) are lean, hyperactive, and have an increased metabolic rate (Kokkotou et al., 2005; Shimada et al., 1998). Similarly, *Mchr1* deletion also results in leanness, hyperactivity, and increased energy expenditure (Åstrand et al., 2004; Chen et al., 2002; Marsh et al., 2002). Although MCH administration stimulates feeding (Qu et al., 1996; Sears et al., 2010), transgenic models of *Mch* or *Mchr1* deletion do not similarly effect feeding (Lord et al., 2022). By contrast, hyperactivity and increased energy expenditure are consistent features following the transgenic *Mch* or *Mchr1* deletion.

A hyperdopaminergic state underlies the hyperactivity of *Mch-* or *Mchr1-KO* animals. There is an elevated amount of dopamine in the accumbens nucleus of *Mch*-*KO* mice (Pissios et al., 2008) and rats (Mul et al., 2011). Furthermore, treating *Mch-KO* mice with a dopamine reuptake inhibitor leads to greater accumulation of dopamine at the accumbens, and this can enhance hyperlocomotor activity (Pissios et al., 2008). Consistently, MCH can suppress dopamine release in the accumbens, and the inhibitory actions of MCH are mediated in part by *Mchr1* expression at GABAergic accumbens neurons (Chee et al., 2019).

Dopaminergic afferents in the accumbens originate from the ventral tegmental area (VTA) (Albanese & Minciacchi, 1983) and form the mesolimbic pathway implicated in the regulation of reward and motivation. The VTA is a heterogeneous area comprising dopaminergic, GABAergic, and glutamatergic cells (Morales & Margolis, 2017), as well as cells that are both GABAergic and glutamatergic (Root, Mejias-Aponte, Qi, et al., 2014; Yamaguchi et al., 2007), dopaminergic and GABAergic (Stamatakis et al., 2013; Tritsch et al., 2012), or dopaminergic and glutamatergic (Dal Bo et al., 2004; Stuber et al., 2010; Sulzer et al., 1998; Tecuapetla et al., 2010). Interestingly, dopaminergic VTA neurons receive inputs from local GABA and glutamate cells (Morales & Margolis, 2017), but other brain regions, such as the lateral hypothalamic area, projecting to the VTA may also regulate dopaminergic VTA neurons (Nieh et al., 2016).

MCH neurons may directly innervate multiple VTA cell types, including GABAergic, glutamatergic, and dopaminergic cells (Faget et al., 2016). Optogenetic activation of MCH neurons can increase dopamine release in the accumbens and increase sucrose preference over a non-nutritive sweetener (Domingos et al., 2013). However, *Mch-KO* mice do not show deficits in nutrient-sensing because the nutritive value associated with sucrose is ascribed to glutamate release from MCH neurons (Schneeberger et al., 2018). Nevertheless, this does not preclude direct MCH actions on VTA neurons that can inhibit dopamine release.

The VTA comprises MCH-immunoreactive fibers (Bittencourt et al., 1992) and *Mchr1* mRNA (Hervieu et al., 2000), but the identity of MCHR1-expressing VTA cells and mechanism(s) of MCH action in the VTA have not been defined. We determined *Mchr1* mRNA expression on GABAergic, glutamatergic, or dopaminergic VTA cells identified by the vesicular GABA transporter (*Vgat*), vesicular glutamate transporter 2 (*Vglut2*), or tyrosine hydroxylase (TH), respectively. We found that MCH directly hyperpolarized dopaminergic or GABAergic VTA cells but not glutamatergic cells. As VTA dopamine cells receive both GABAergic and glutamatergic afferents (Marisela Morales & Margolis, 2017), we also determined if MCH may regulate GABAergic or glutamatergic synaptic transmission at dopaminergic VTA cells. Interestingly, MCH did not alter GABAergic transmission at dopaminergic VTA cells, but it produced a delayed increase in glutamatergic input. This may reflect a disinhibition of glutamatergic tone onto dopamine cells in the VTA, as MCH can suppress GABAergic transmission at glutamate cells. These findings supported both direct and indirect MCH action that regulates neuronal activity or synaptic transmission at dopaminergic VTA cells.

## Materials and Methods

All animal procedures were approved by the Animal Care Committee at Carleton University. All mice were bred on a C57BL/6J background and group housed in a 12:12 hour light:dark cycle with *ad libitum* access to water and standard mouse chow (Teklad Global Diets 2014, Envigo, Mississauga, Canada).

## Quantitative real-time PCR

WT mice 6–20 weeks old (N = 16) were anesthetized with an intraperitoneal injection (ip) of 7% chloral hydrate (700 mg/kg). The brain was rapidly removed from the skull and samples of the striatum, hypothalamus, hippocampus, cerebellum, and VTA were microdissected and flash-frozen in liquid nitrogen. Tissue samples were homogenized in TRIzol (Invitrogen, Carlsbad, CA), and RNA was isolated by chloroform and precipitated by isopropanol. The RNA pellet was washed with 70% ethanol and air-dried before resuspending in distilled water. RNA purity was determined by a 260/280 absorbance ratio of ∼2.0 using a NanoDrop Lite Spectrophotometer (Thermo Scientific, Waltham, MA).

RNA samples were treated with DNase and reverse transcribed for cDNA synthesis with an iScript gDNA Clear cDNA Synthesis Kit (1725035, Bio-Rad, Hercules, CA). We performed quantitative RT-PCR using a CFX Connect Real-Time PCR Detection System thermal cycler (1855201, Bio-Rad). Custom primers (Invitrogen) for mouse glyceraldehyde-3-phosphate dehydrogenase (*Gapdh*) and *Mchr1* mRNA comprised the following sequences: *Gapdh* forward, 5’-ATGTGTCCGTCGTGGATCTGA-3’; *Gapdh* reverse, 5’-ATGCCTGCTTCACCACCTTCTT-3’; *Mchr1* forward, 5’-CAATGCCAGCAACATCTCC-3’; *Mchr1* reverse, 5’-ACCAAACACTGAAGGCATGA-3’.

*Mchr1* mRNA expression in each brain region was calculated via double delta Ct analysis and standardized to the expression levels of *Gapdh* of the same mouse. Relative *Mchr1* mRNA expression in each brain region was then normalized to the level of *Mchr1* mRNA in the hypothalamus for each sex.

## Histology

### Tissue preparation

Mice (12–20 weeks old) were anaesthetized with 7% chloral hydrate (700 mg/kg, ip) and underwent transcardiac perfusion with cold (4 °C) sterile saline (0.9% NaCl) followed by pre-chilled (4°C) phosphate buffered formalin (10%; VWR, Radnor, PA).

### Immunohistochemistry (IHC)

Brain tissue from WT (N = 6) and *Mchr1*-*KO* mice (N = 2) were post-fixed in 20% sucrose prepared in formalin (4 h) and then cryoprotected with 20% sucrose prepared in phosphate buffered saline (PBS, pH 7.4) for 16 h until sectioning using a freezing microtome (Spencer Lens Co., Buffalo, NY) into five series of 30 μm coronal sections. MCHR1 immunoreactivity was labeled as previously described (Diniz et al., 2020). In brief, brain tissue was treated with 0.3% hydrogen peroxide and incubated with a rabbit anti-MCHR1 antibody (1:1,000; PA5-24182, UC2738292, Invitrogen, Carlsbad, CA) for 48 h at 4 °C in a 3% normal donkey serum (NDS) solution. Tissue was thoroughly rinsed, incubated with a biotinylated donkey anti-rabbit IgG antibody (1:500; 711-065-152, Lot 101909, Jackson ImmunoResearch Laboratories) at RT for 1 h, and treated with avidin-biotin-horseradish peroxidase (1:1:833; PK-6100, Vector Labratories, Burlingame, CA) at RT for 30 min before incubating with 0.5% biotinylated (EZ-Link sulfo-NHS-LC-biotin; ThermoFisher Scientific, Waltham, MA) tyramine (Sigma-Aldrich; Sigma Chemical; St. Louis, MO, USA) for 20 min (RT). All sections were then incubated with an Alexa Fluor 647-conjugated streptavidin (1:500; 016-600-084, Lot 135095, Jackson ImmunoResearch Laboratories) and NeuroTrace (1:50; N21479, ThermoFisher) for 2 h at RT. Following staining, all tissue was thoroughly rinsed and immediately mounted onto Superfrost Plus microscope glass slides (ThermoFisher) and coverslipped with Prolong Diamond Antifade mounting media (P36961, Invitrogen).

After IHC processing for MCHR1 immunoreactivity, one additional series were incubated with a sheep anti-TH antibody (1:1,000; GTX82570, Lot 822005648, GeneTex, Irvine, CA) at RT overnight. Sections were then incubated with a donkey anti-sheep secondary antibody conjugated with Alexa Fluor 594 (1:500; 715-496-150, Lot 82079, JacksonImmuno Research Laboratories) and NeuroTrace (1:50) for 2 h at RT. Tissues were rinsed, immediately mounted, and coverslipped as described above.

### Thick tissue IHC

Brain slices following patch-clamp recordings were fixed in formalin for at least 24 h, treated with a 2% Triton-X PBS solution for 45 min at RT to permeabilize the tissue, and then incubated with a sheep anti-TH primary antibody (1:1,000; GeneTex) at 4 °C for 72 h. After rinsing, the tissue was incubated with a streptavidin-conjugated Cy3 (1:500; 016-160-084, Lot 146014, JacksonImmuno Research Laboratories) and donkey anti-sheep conjugated with Cy5 (1:500; 713-176-147, Lot 70443, JacksonImmuno Research Laboratories) secondary antibodies. The tissues were then rinsed and immediately mounted onto Superfrost Plus microscope glass slides and coverslipped with Prolong Diamond Antifade mounting media).

### Combined *in situ* hybridization (ISH) and IHC

Brains from WT mice (N = 6) were post-fixed in 10% formalin overnight, cryoprotected in 20% sucrose with 0.5% sodium-azide (24 h) and sectioned using a freezing microtome into five series of 30 μm coronal sections. The series designated for ISH were immediately mounted onto Superfrost Plus microscope slides, air-dried at room temperature (RT; 21–23 °C; 1 h) and −20 °C (30 min), and then stored at −80 °C. Adjacent brain series were reserved for Nissl staining, RNAscope probe labeling, and negative control, respectively. One slide from a separate series served as the positive control. We determined the coexpression of *Mchr1* with *Slc32a1* (*Vgat*) and *Slc17a6* (*Vglut2*) by RNAscope-enabled ISH and with TH by subsequent IHC in the same brain tissue. ISH procedures were completed based on manufacturer instructions for RNAscope ISH on fixed-frozen mouse brain tissue (Multiplex Fluorescent Reagent Kit v2 User Manual; Advanced Cell Diagnostics (ACD), Newark, CA) as previously described (Bono et al., 2022), unless indicated otherwise.

In brief, slide-mounted brain tissues were dehydrated in an ethanol gradient, treated with hydrogen peroxide and target retrieval procedure for 5 min (99 °C), and then dehydrated and air-dried overnight (RT). Tissues of interest were isolated by drawing a hydrophobic barrier and treated with Protease III in a HybEZ oven (40 °C, 30 min). Slides were then incubated with a cocktail of *Mm-Mchr1* (317491, ACD) in channel 1 (C1), *Mm-Slc32a1-C2* in C2 (319191-C2, ACD), and *Mm-Slc17a6* in C3 (319171-C3, ACD) probes (40 °C, 2 h). Concentrated *Mm-Slc32a1* C2 and *Mm-Slc17a6* C3 probes were added to the diluted *Mm-Mchr1* C1 probe at a ratio of 1:50. A volume of 5–8 drops of the probe mixture was added to each slide. *Bacilus* dihydrodipicolinate reductase (*dapB*, 320871, ACD) and *Mm-Ppib* (313911, ACD) probes were used as the negative and positive control, respectively. Hybridization signals were amplified by sequential incubation at 40 °C with AMP-1 (30 min; 323101, ACD), AMP-2 (30 min; 323102, ACD), and AMP-3 (15 min; 323103, ACD).

C1 *Mchr1* hybridization was developed (40 °C, 15 min with HRP-C1; 323104, ACD) and tyramide signal amplification (TSA) plus Opal 690 (1:750, 40 °C, 30 min; FP1497001KT, Akoya Biosciences, Marlborough, MA), then quenched with HRP Blocker (40 °C, 15 min; 323107, ACD). C2 *Vgat* and C3 *Vglut2* hybridization were sequentially developed with HRP-C2 (323105, ACD) with TSA plus Cyanine 3 (1:1,500; NEL44E001KT, PerkinElmer, Waltham, MA) and HRP-C3 (323106, ACD) with TSA plus Opal 520 (1:750; FP1487001KT, Akoya Biosciences). C2 signal was similarly quenched with HRP Blocker (40 °C, 15 min) before proceeding to C3 development.

Following ISH of probe tissue, the tissues were treated for IHC labeling of TH immunoreactivity based on ACD directions for combined ISH and IHC (Advanced Cell Diagnostics, 2020) and as previously described (Mickelsen et al., 2019). Tissues were blocked with 3% NDS solution prepared in PBS (700 μl) for 30 min (RT) and then incubated with anti-sheep TH primary antibody (1:1,000; GTX82570, Lot 822005648, GeneTex, Irvine, CA) 2 h (RT). Slides were thoroughly rinsed in PBS and then incubated with a donkey anti-sheep secondary antibody conjugated with Alexa Fluor 594 (1:500; 715-496-150, Lot 82079, JacksonImmuno Research) for 30 min (RT). Slides were rinsed in Wash Buffer (320058, ACD), treated with DAPI Reagent (323108, ACD) for 30 s (RT), and immediately coverslipped with ProLong Gold Antifade Mountant (Fisher Scientific). Slides were kept in a dark drying drawer at RT overnight and then transferred to −20 °C for storage until imaging.

### Nissl stain

Sections underwent Nissl staining as previously described (Bono et al., 2022; Negishi et al., 2020).

## Microscopy

### Brightfield imaging

Nissl-stained tissue was acquired with a Nikon Eclipse Ti2 inverted microscope (Nilon Instruments Inc., Mississauga, Canada) equipped with a motorized stage and DS-Ri2 color camera (Nikon) using a Plan Apochromat 10x objective (0.45 NA). Brightfield images were stitched at 20% overlap with NIS Elements software.

### Confocal imaging

Images from Nissl-stained tissue were aligned to tiled, overview confocal images showing the VTA in relation to other midbrain regions acquired using the 405-nm wavelength laser with a 4x Plan Apochromat objective (0.2 NA). Stitched, high resolution confocal frames from IHC- and/or ISH-treated tissues were captured from within the overview image.

***Confocal imaging of MCHR1 immunoreactivity*** underwent scans with a 10x and/or 20x (0.75 NA) Plan Apochromat objectives using a Nikon C2 confocal system. Images were acquired with a 405- and 647-nm wavelength laser to visualize NeuroTrace- and Alexa Fluor 647-labeled signals. Tissue that was also stained for TH had additional images acquired with a 561-nm wavelength laser to visualize Alexa Fluor 594-labeled signal, and fluorescence signals from the 561- and 647-nm channels were pseudocoloured magenta and green, respectively.

***Confocal imaging of combined ISH- and IHC-treated tissue*** underwent scans with 20x Plan Apochromat objective using a Nikon C2 confocal system and acquired with a 405-, 488-, and 647-nm wavelength laser to visualize DAPI-, Opal 520-, and Alexa Fluor 647-labeled signals for nuclear stains, *Vglut2* hybridization, and *Mchr1* hybridization, respectively. Simultaneously at the same optical section, Cyanine 3- and Alexa Fluor 594-labeled signals to visualize *Vgat* hybridization and TH were acquired by spectral imaging using a 561-nm wavelength excitation laser in the continuous band (CB) setting to generate a lambda stack acquired with sequential 10-nm bandwidths spanning the wavelength range of 580 nm to 640 nm. The spectral profiles were unmixed to separate Cyanine 3 and Alexa Fluor 594 signals, which were most distinguished at 590 nm and 620 nm, respectively, and then merged with signals acquired by the 405-, 488-, and 647-nm wavelength laser to generate one image file displaying signals from all five fluorophores.

Tissues that served as the negative control were also acquired using the same combination of conventional confocal and spectral unmixing for probe tissue. Background fluorescence related to nonspecific binding from ISH treatment was determined using the negative control tissue by adjusting the look up table (LUT) values for each of the three image channels to yield a resultant black image. The LUT range was then applied to images acquired from the probe tissue to subtract background fluorescence, thus any remaining signal observed from the probe tissue was considered to be positive hybridization signals.

***Confocal imaging of IHC-treated cells in thick tissue*** underwent scans with a 20x Plan Apochromat objective in a Nikon C2 confocal system and acquired with a 488-, 561-, and 647-nm wavelength laser to visualize native EGFP fluorescence, Cy3-labelled signal for biocytin-filled cell, and Cy5-labelled signals for TH immunoreactivity, respectively. Cy3 signal was pseudocoloured blue.

## Neuroanatomical analyses

### Plane-of-section analysis

Nissl or fluorescent-Nissl images were parcellated in Adobe Illustrator CS6 (Adobe Inc., San Jose, CA) based on the regions defined by the *Allen Reference Atlas* (*ARA*; Dong, 2008), as previously described (Bono et al., 2022; Negishi et al., 2020). Parcellations for MCHR1-ir cells were directly applied from fluorescent-Nissl signals captured within the same image acquisition. High magnification confocal images of ISH-treated tissues were indirectly parcellated from brightfield images of an adjacent Nissl-stained section, which was aligned to the low magnification, overview confocal image of the probe tissue.

### Identifying MCHR1- and TH-ir signals

Our MCHR1 antibody (1:1,000; PA5-24182, Invitrogen, Carlsbad, CA) labeled MCHR1 immunoreactivity at the primary cilium of neurons (Diniz et al., 2020). Only cilia, identified by their long spindle-like appearance greater than 1 μm (Bansal et al., 2021; Diniz et al., 2020), associated with a NeuroTrace-labeled somata was counted as a MCHR1-expressing cell. Only TH staining at the somata that colocalized with DAPI-, NeuroTrace-, or EGFP-labeled cell was considered. Only signals within the boundaries of the VTA, as determined by our plane-of-section analysis, were included in our analyses.

### Identifying hybridization signals

Signal criteria was independently-established based on the signal-to-background level for each hybridization signal. *Mchr1* hybridization signal appeared as punctate “dots” with clusters centered tightly around a DAPI-labelled nucleus (Bono et al., 2022). *Vgat* hybridization appeared as dot clusters that took the shape of cell bodies and were colocalized to a DAPI-labelled nucleus. Compared to *Vgat* hybridization, dot clusters following *Vglut2* hybridization was less dense within the cell and may appear as punctate dots.

### Mapping

A circle was placed over each positively-labeled immunoreactive or hybridized cell, and the relative positions of the circles were transferred, as previously described (Bono et al., 2022; Negishi et al., 2020), to coronal brain atlas templates (Dong, 2008) to generate standardized maps of *Mchr1*, *Vglut2*, *Vgat*, and TH expression. Cell counts at each atlas level were determined from cells mapped and counted unilaterally.

Number of cells reported for IHC and combined ISH and IHC studies were corrected for oversampling, as previously described (Chee et al., 2013; Negishi et al., 2020; Bono et al., 2022), using the Abercrombie formula 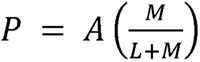 (Abercrombie, 1946): where P is the corrected count, A is the original count, M is the mean tissue thickness (IHC, 13.6 μm; ISH+IHC, 10.9 μm) determined from five brain slices, and L is the mean cell diameter (IHC, 15.6 μm; ISH+IHC, 16.0 μm) determined from 80 cells.

## Electrophysiology

### Animals

*Th-cre;L10-Egfp*, *Vgat-cre; L10-Egfp*, and *Vglut2-cre;L10-Egfp* reporter mice were generated by crossing *Th-cre* (Lindeberg et al., 2004), *Vgat-cre* (028862, JAX; Vong et al., 2011), or *Vglut2-cre* (028863, JAX; Vong et al., 2011) mice, respectively to homozygous *lox-STOP-lox-L10-Egfp* reporter mice (Krashes et al., 2014). All transgenic lines were kindly provided by Dr. B. B. Lowell (Beth Israel Deaconess Medical Center, Boston, MA).

### Acute slice preparation

Mice (4–10 weeks old) were anaesthetized with 7% chloral hydrate (700 mg/kg, ip) and transcardially perfused with ice cold, carbogenated (95% O_2_, 5% CO_2_) slice artificial cerebrospinal fluid (ACSF) solution (300 mOsm/L; pH 7.4) containing (in mM) 92 NMDG, 2.5 KCl, 1.25 NaH_2_PO_4_, 30 NaHCO_3_, 20 HEPES, 25 glucose, 2 thiourea, 5 Na-ascorbate, 3 Na-pyruvate, 0.5 CaCl_2_·2H_2_O, and 10 MgSO_4_ (Ting et al., 2018). Brains were sectioned on a vibratome (VT 1000S, Leica Biosystems, Wetzlar, Germany) in slice ACSF to produce 250 μm-thick coronal slices that were immediately transferred to carbogenated slice solution at 37 °C for 5 min. Slices were then incubated in a carbogenated bath ACSF (300 mOsm/L) containing (in mM) 124 NaCl, 3 KCl, 1.3 MgSO_4_, 1.4 NaH_2_PO_4_, 10 D-glucose, 26 NaHCO_3_, and 2.5 CaCl_2_ for 5 min and then allowed to recover in bath ACSF (RT) for at least 1 hr prior to recordings.

### Patch-clamp recordings

Brain slices containing the VTA, which primarily encompassed L83–87, were placed in a slice chamber, superfused (2–2.5 ml/min) with carbogenated bath ACSF pre-warmed (37 °C) by a temperature controller (Warner Instrument Corporation, Hamden, CT), and held in place by a platinum ring strung with nylon fiber made in-house.

VTA cells were visualized with infrared differential interference contrast microscopy at 40x magnification on either an Examiner.A1 microscope (Zeiss, Oberkochen, Germany) equipped with an AxioCam camera (Zeiss) and Axiovision software (Zeiss) or with an Eclipse FN1 microscope (Nikon) equipped with a pco.panda 4.2 camera (Excelitas PCO GmbH, Kelheim, Germany) and NIS-Elements Imaging software (Nikon). EGFP-labeled cells were identified using epifluorescence illumination from a halogen (HXP120, Zeiss) or mercury lamp (C-SHG1, Nikon).

Borosilicate glass pipettes (593800; A-M Systems) were pulled on a P-1000 Flaming/Brown micropipette puller (Sutter Instruments; Novato, CA) and backfilled (7–10 MΩ) with a potassium-based internal solution (290 mOsm/L, pH 7.24) containing (in mM) 120 K-gluconate, 10 KCl, 10 HEPES, 1 MgCl_2_, 1 EGTA, 4 MgATP, 0.5 NaGTP, and 10 phosphocreatine to assess membrane properties and glutamatergic events. A cesium-based internal solution (290 mOsm/L, pH 7.24) containing (in mM) 128 CsMS, 11 KCl, 10 HEPES, 0.1 CaCl2, 1 EGTA, 4 MgATP, 0.5 NaGTP) was used to record GABAergic events. Voltage-clamp recordings were acquired at a holding potential of V_h_ = −60 mV and −5 mV for excitatory and inhibitory events, respectively using a MultiClamp 700B amplifier (Molecular Devices, San Jose, CA) and digitized using a Digidata 1440A (Molecular Devices). Biocytin (0.4%; Cayman Chemical, Ann Arbor, MI) was added to the internal solution for post-hoc thick tissue IHC in a subset of recorded cells. Traces were acquired at 10 kHz and filtered at 3 kHz by pClamp 10.3 or 11 software (Molecular Devices), as previously described (M. Chee et al., 2010).

### Drug application

Pharmacological applications of MCH (H-1482, Bachem, Torrance, CA) was applied by bath perfusion or puff application after a baseline period of at least 5 min or 1 min, respectively, where the membrane potential fluctuated by less than 10%. Cells were held for at least 20 min during the MCH-washout period to assess the reversibility of MCH-mediated effects on the membrane potential. Where applicable, tetrodotoxin citrate (TTX; 500 nM; T-550, Alomone labs, Jerusalem, Israel) or the MCHR1-antagonist TC-MCH 7c (10 μM; 4365, Tocris, Toronto, Ontario, Canada), was bath applied and allowed to saturate the slice for at least 10 min prior to MCH application and then maintained during the MCH application and washout period as well. Puff-applied MCH was delivered via a puff-pipette backfilled with 3 μM MCH (or bath ACSF) positioned within 20–40 μm away from the patched cell held in the whole-cell configuration. A short “puff” was applied by manually applying positive pressure via the puff-pipette, and a solution ripple can be seen to contact the patched cell.

## Experimental design and statistical analyses

### qPCR studies

Comparison of *Mchr1* gene expression in wildtype mice (N = 6–7/sex) were analyzed using a between-subject design. *Mchr1* expression in different regions were analyzed using a two-way mixed model ANOVA with Šidák post-hoc testing to determine sex differences.

### Neuroanatomical studies

Comparisons of the total number of MCHR1-expressing cells were analyzed using a between-subject design from (N = 3/sex) and compared via unpaired *t* test, and differences in the distribution of MCHR1-expressing cells across each *ARA* level between sexes was determined by the two-way mixed model ANOVA.

Comparisons of *Mchr1*-, *Vglut2*-, *Vgat*-, or TH-expressing cells were analyzed from male and female wildtype mice (N = 6) using a between-subject design, and a one-way ANOVA was used to identify distinct VTA cell groups. Sex differences were determined by two-way ANOVA.

### Electrophysiological studies

Acute brain slices were prepared from male and female *Th-cre;L10-Egfp* (N = 27), *Vgat-cre;L10-Egfp* (N = 8), and *Vglut2-cre;L10-Egfp* (N = 8) reporter mice. Up to three cells were recorded from three to four coronal slices, where the VTA sits between the medial mammillary nucleus and compact part of the substantia nigra in rostral sections and sits beside the interpeduncular nucleus in caudal sections. Recordings from EGFP-labeled VTA cells of each reporter line was guided by atlas maps of *Mchr1* expression in each cell type. Electrical signals recorded were processed in Clampfit 10.7 (Molecular Devices).

The resting membrane potential (RMP) was sampled every 1 s and averaged into 30-s bins in bath-applied treatments and sampled every 100 ms and averaged into 1-s bins in puff-applied treatments. Cells deemed to be MCH-responsive exhibited a change in RMP lasting at least 2 min that was greater than 2-times the standard deviation of the mean RMP at baseline. MCH-mediated change in RMP (Δ RMP) was sampled within 7 min after MCH application or within 5 s of puff application. Comparisons of mean RMP before and after MCH application was determined via a paired *t* test and reported with effect size (η^2^). Comparisons of MCH-mediated Δ RMP over time in the presence or absence of TTX was determined using within-subject design and repeated measures two-way ANOVA with Šidák post-hoc testing.

Excitatory (EPSC) and inhibitory postsynaptic current (IPSC) events were detected using MiniAnalysis (Synaptosoft, Decatur, GA). Event detection threshold was set to 5-times the root mean square noise level for each recording. The frequency of EPSCs and IPSCs were indirectly determined by the interevent interval between events using MiniAnalysis (Synaptosoft, Decatur, GA) and averaged into 30-s bins. The baseline event frequency or amplitude was determined as the mean event frequency averaged over 5 min immediately prior to MCH bath application. Comparisons of mean event frequency or amplitude before and after MCH application was determined via a paired *t* test and reported with effect size (η^2^). Comparisons of MCH-mediated effects on postsynaptic current events over time in the presence or absence of TTX or TC-MCH 7c was determined using within-subject design and repeated measures two-way ANOVA with Šidák post-hoc testing.

### Graphs

Only responders were included in our graphical representations, unless a null response was referenced. All data graphs were prepared using Prism 9 (GraphPad, San Diego, CA). Results were considered statistically significant at p < 0.05. All data are reported as group mean ± SEM. Figures were assembled in Adobe Illustrator 2020 (Adobe Inc.). Numeral in parentheses in graphs represent the number of animals (N) or cells (n).

## Results

### Moderate *Mchr1* mRNA and MCHR1 protein expression in the mouse VTA

*Mchr1* mRNA and protein have been reported in the rat VTA (Hervieu et al., 2000; Saito et al., 2001), and we first determined *Mchr1* mRNA and/or protein expression in the VTA of male and female wildtype mice. While the level of *Mchr1* mRNA varied across brain regions (F(4, 44) = 81.6, p < 0.0001, ANOVA), it was similar between male and female mice (F(1, 11) = 0.6, p = 0.449, ANOVA), and there was no interaction between brain region and sex (F(4, 44) = 0.8, p = 0.561, ANOVA). As expected, *Mchr1* expression was highest in the striatum, which included the nucleus accumbens and caudate putamen, and barely detectable in the cerebellum (**Figure 1A**). Importantly, *Mchr1* expression in the VTA was comparable to the hypothalamus (male: t(44) = 1.5, p = 1.000; female: t(44) = 1.0, p = 1.000) and hippocampus (male: t(44) = 1.3, p = 1.000; female: t(44) = 0.6, p = 1.000; **Figure 1A**).

**Figure 1.**
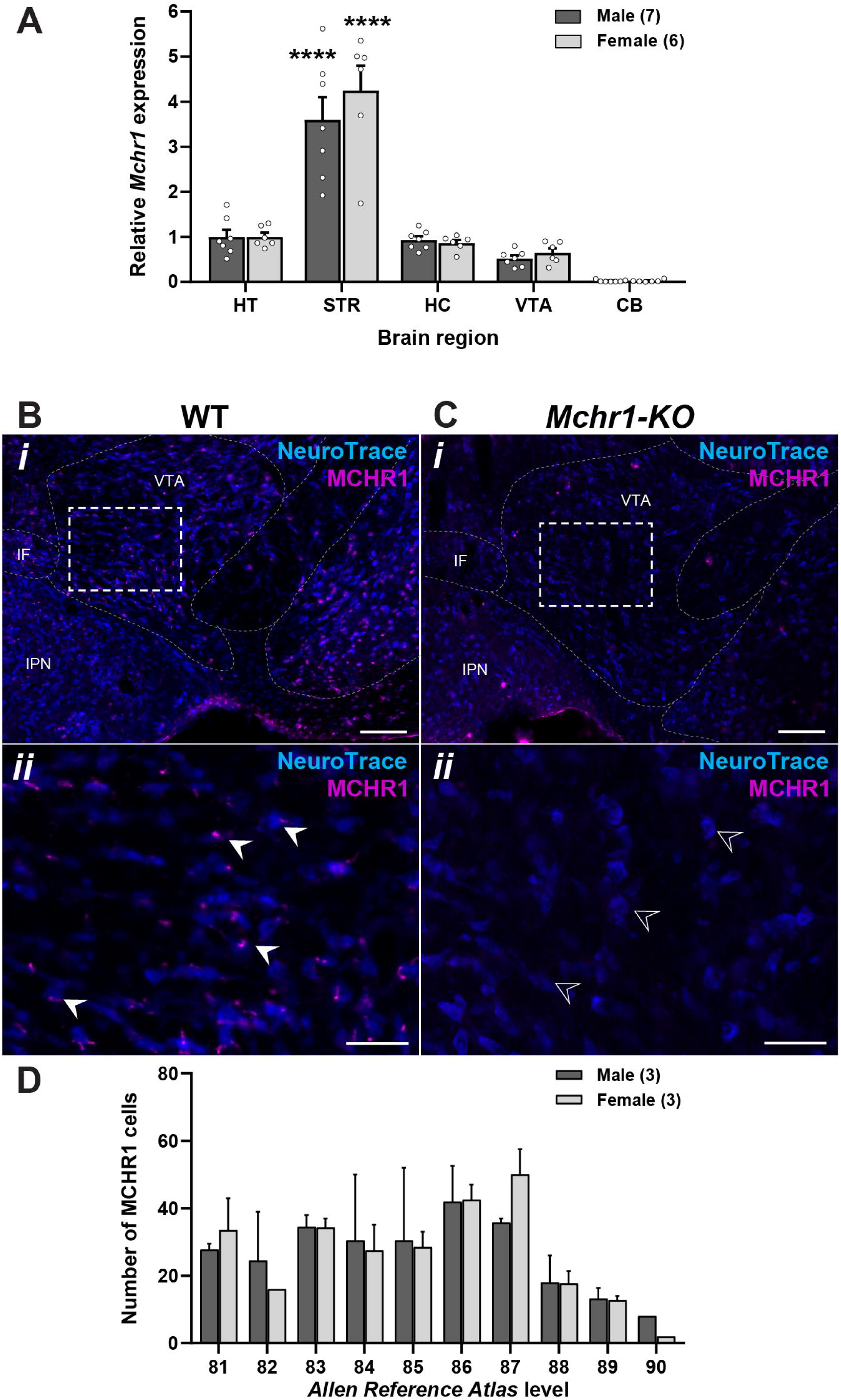
*Mchr1* mRNA and MCHR1 protein expression in wildtype mouse VTA. qPCR analysis of *Mchr1* mRNA from the hypothalamus (HT), striatum (STR), hippocampus (HC), ventral tegmental area (VTA), and cerebellum (CB) of male and female wildtype mice (**A**). Confocal photomicrographs of MCHR1-immunoreactive signals on Nissl-stained (NeuroTrace) soma in wildtype (**B**) and *Mchr1-KO* (**C**) VTA at low (***i***) and high magnification (***ii***, from dash-outlined region in ***i***). Representative examples of the presence (white arrowhead) or absence (open arrowhead) of MCHR1 immunoreactivity at the primary cilium of VTA cells. Number of MCHR1-immunoreactive cells throughout the VTA of male and female wildtype mice (**D**). VTA levels were assigned with reference to the *Allen Reference Atlas* (*ARA*; Dong, 2008). Two-way ANOVA with Bonferroni post-test: ****, p < 0.0001 (STR vs all regions; **A**). Scale bars: ***i***, 100 µm; ***ii***, 40 µm. IF, interfascicular nucleus raphe; IPN, interpeduncular nucleus; VTA, ventral tegmental area.

The VTA of wildtype male and female mice also comprised MCHR1-immunoreactive (MCHR1-ir) signals, which appeared long and spindle-like and were consistent with MCHR1 expression observed on the primary cilium of neurons (**Figure 1B**). No MCHR1-ir labeling was detected in the VTA of *Mchr1-KO* mice (**Figure 1C**). Male and female wildtype mice had a similar distribution pattern of MCHR1-expressing cells throughout the VTA (F(1, 4) = 0.02, p = 0.891, ANOVA; **Figure 1D**). Additionally, there was no difference in the total number of MCHR1-expressing cells in the VTA of male (159 ± 48, n = 3) and female (171 ± 29, n = 3) mice (t(4) = 0.2, p = 0.835, η^2^ = 0.01, unpaired *t* test).

## Neurochemical diversity of VTA cells

The VTA comprises multiple cell types, including dopaminergic, GABAergic, or glutamatergic cells that have distinct distribution patterns (M. Morales & Root, 2014; Marisela Morales & Margolis, 2017). To identify the neurochemical identities of VTA cell types, we determined if *Mchr1* hybridization colocalized with *Vglut2* or *Vgat* hybridization in glutamatergic or GABAergic cells, respectively, or with TH immunoreactivity in dopaminergic cells within the same tissue. As there were no sex differences in *Mchr1* mRNA (**Figure 1A**) or protein expression (**Figure 1D**), we herein combined our neuroanatomical analyses from male and female mice.

The overall number of VTA cells labeled by *Vglut2* mRNA, *Vgat* mRNA, and/or TH immunoreactivity decreased posteriorly (**Figure 2A**, gray line). We compared the distribution of *Vglut2* mRNA, *Vgat* mRNA, and TH-ir cells within the VTA to determine their relative prevalence rostrocaudally, and we distinguished at least five subtypes of VTA cells (F(4, 50) = 13.4, p = 0.0001, ANOVA). *Vglut2* cells were most prominent anteriorly then gradually diminished towards the posterior VTA (**Figure 2A**), and they were usually medially distributed within the VTA (**Figure 2B**). *Vgat* cells comprised ∼30% of VTA cells counted through the anterior and middle sections of the VTA (L80–87) but made up the majority of cells in the posterior VTA (L88–90) (**Figure 2A**). TH-ir cells were most numerous in the middle of the VTA and comprised ∼40% of cells counted (**Figure 2A**). Both *Vgat* and TH cells tended to be laterally distributed, though some *Vgat* cells were spread medially (**Figure 2B**).

**Figure 2.**
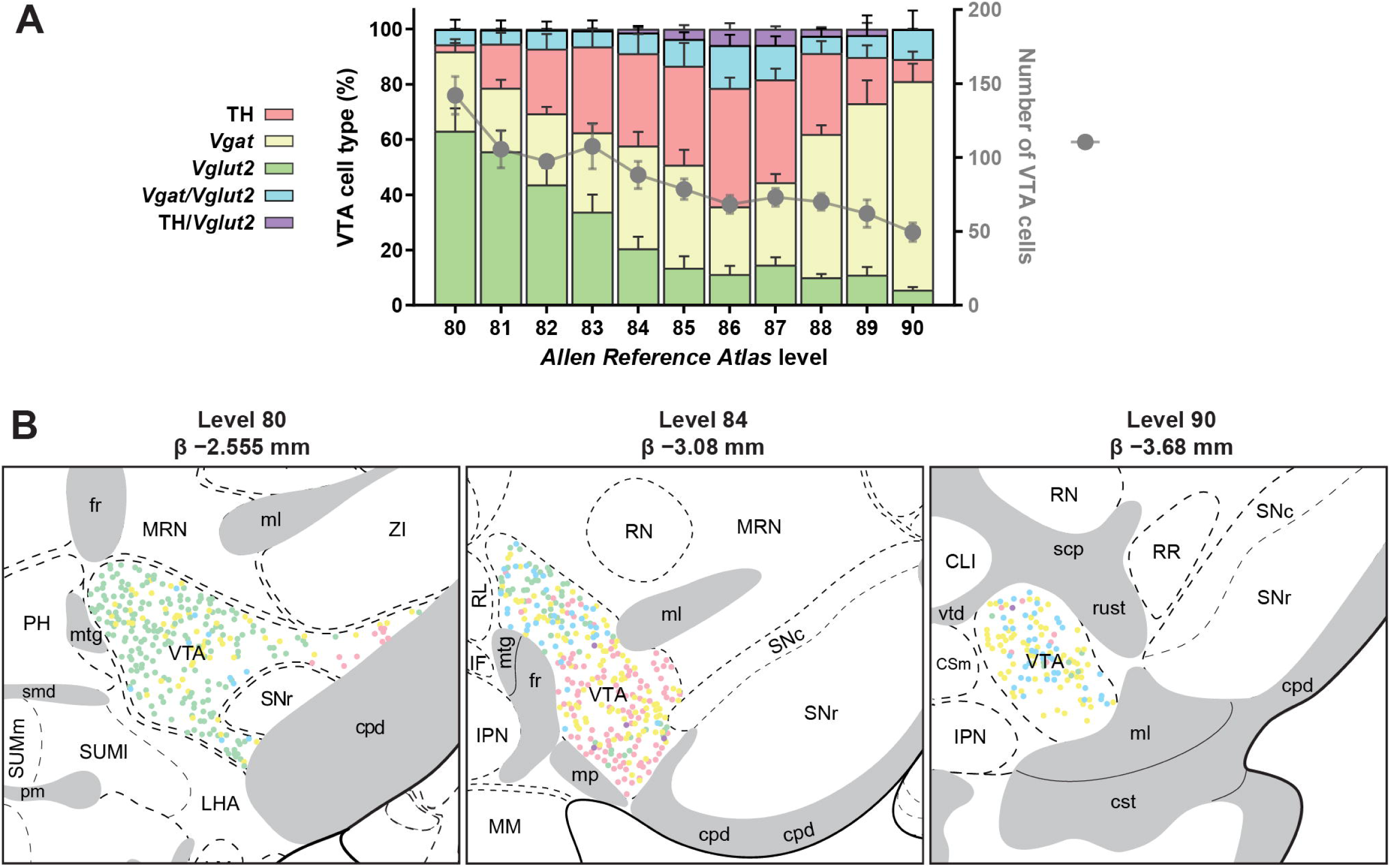
Neurochemical characterization of VTA cells. Number (right *y*-axis, gray line) and proportion of each cell type (left *y*-axis, stacked colored bars) at each *Allen Reference Atlas* level (Dong, 2008) rostrocaudally from VTA slices expressing *Vglut2* mRNA hybridization, *Vgat* mRNA hybridization, and tyrosine hydroxylase (TH) immunoreactivity (N = 6 mice; **A**). Representative coronal brain maps of VTA cells that are *Vglut2*-, *Vgat*- and/or TH-positive (**B**). Each panel includes the nomenclature, atlas level and Bregma value (β) assigned to that tissue in accordance with the *Allen Reference Atlas* (Dong, 2008).

In addition, we also found that VTA cells may express two or more neurotransmitter markers. The most common were those expressing both *Vglut2* and *Vgat* that comprised 5–10% of VTA cells and up to 16% of cells towards the middle of the VTA (L86–87; **Figure 2A**). Cells expressing TH and *Vglut2* emerged towards the posterior VTA (L85–89) but comprised <6% at its peak (**Figure 2A**) so were not mapped. Similarly, just 0.25% of cells exhibited TH and *Vgat*, and cells exhibiting *Vglut2*, *Vgat*, and TH were even rarer (<0.05%; data not shown).

## Neurochemical and spatial distribution of *Mchr1* mRNA expression in the VTA

*Mchr1* mRNA expression was widely distributed throughout the VTA (**Figure 3A**) and across all major VTA cell types (**Figure 3B–E**), but there was no sex difference in the heterogeneity of *Mchr1* VTA cells (F(1, 24) = 0.01, p = 0.924, ANOVA; **Figure 3-1A**). Interestingly, about one in four (25%) *Mchr1* cells (*Mchr1*-only) do not coexpress TH, *Vglut2*, or *Vgat* and are yet to be defined neurochemically (**Figure 3F**). These *Mchr1*-only cells can be seen throughout the VTA (**Figure 3G**) and tend to be less prominent in the VTA of female mice, as sex may interact with *Vgat* expression in *Mchr1* cells (F(1, 8) = 4.8, p = 0.059, ANOVA; **Figure 3-1B**). Consistent with the rostrocaudal composition of VTA cells, *Mchr1* expression was predominantly in *Vglut2* cells rostrally, TH cells in the middle sections of the VTA, and in *Vgat* cells caudally (**Figure 3G**).

**Figure 3.**
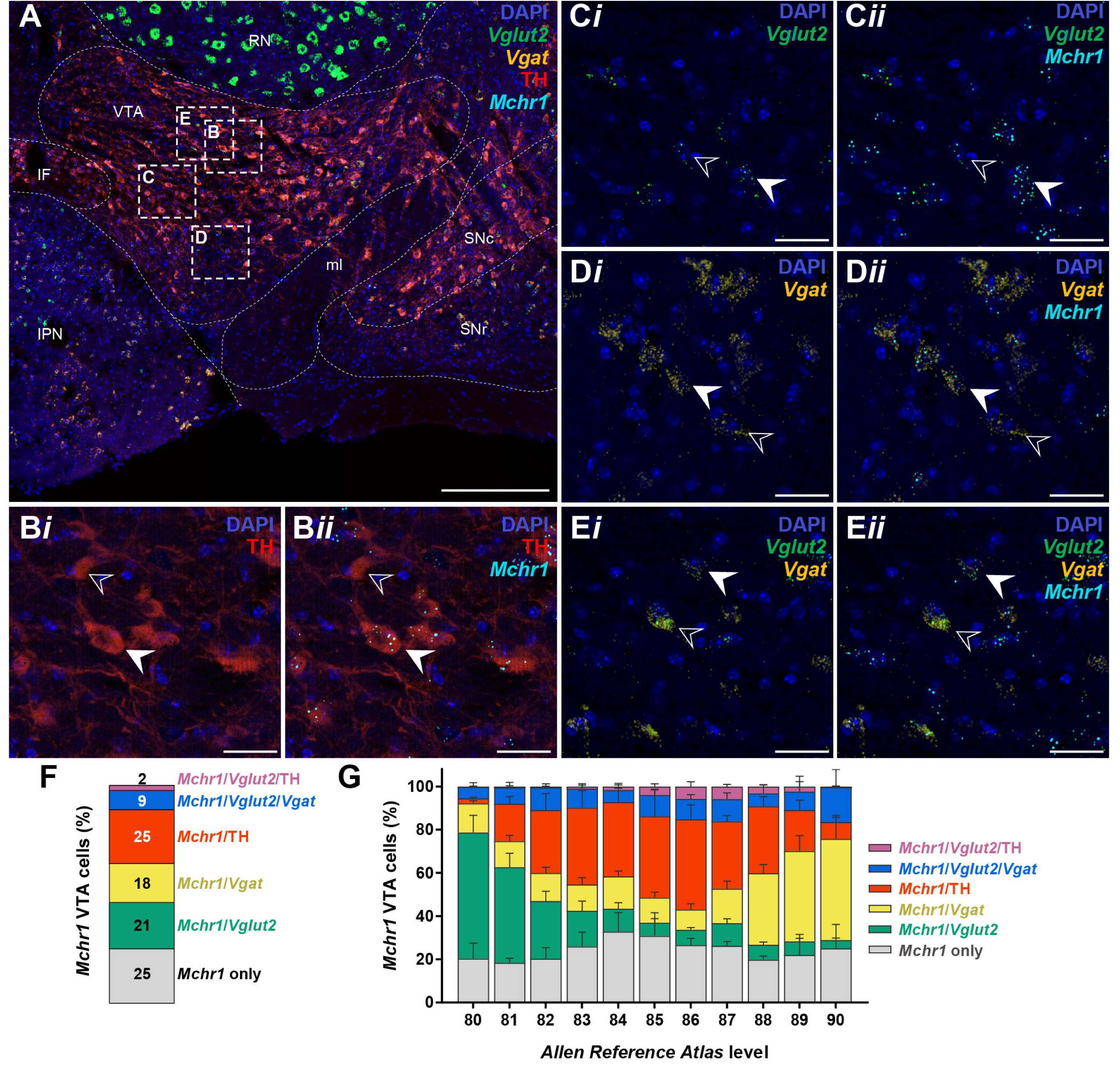
*Mchr1* expression on diverse VTA cell types. Representative low (**A**) and high magnification confocal photomicrographs (**B**–**E**), from corresponding dash-outlined region in **A**) of *Vglut2* mRNA, *Vgat* mRNA, and/or TH immunoreactivity (***i***) that coexpress (white arrowhead) or lack (open arrowhead) *Mchr1* mRNA hybridization in VTA cells (***ii***). Relative proportion of *Mchr1*-expressing VTA cells overall (**F**) and at each level, as determined with reference to the *Allen Reference Atlas* (Dong, 2008), of the VTA (N = 6; **G**). Scale bars: **A**, 250 µm; **B**–**E**, 30 µm. IF, interfascicular nucleus raphe; IPN, interpeduncular nucleus; ml, medial lemniscus; RN red nucleus; SNc, substantia nigra, compact part; SNr, substantia nigra, reticular part; VTA, ventral tegmental area.

About half (52%) of *Vglut2* VTA cells expressed *Mchr1* mRNA (**Figure 4-1**), which was evenly distributed throughout *Vglut2* cells in the medial and anterior VTA but only sparsely distributed posteriorly (**Figure 4A**). By contrast, few *Vgat* cells in the anterior VTA expressed *Mchr1* mRNA (**Figure 4-2**), but in the middle and posterior VTA sections where *Mchr1*-expressing *Vgat* cells were more common, they tend to be laterally distributed or have no discernible pattern, respectively (**Figure 4B**). Interestingly, the majority of TH-ir VTA neurons (78%) expressed *Mchr1* mRNA (**Figure 4-3**), and while *Mchr1*-expressing TH cells were found sparingly in the anterior VTA, they were abundant in middle VTA sections, especially near the ventral and lateral borders of the VTA (**Figure 4C**). In addition, although dual *Vglut2* and *Vgat* cells comprised just one-tenth of all VTA cells, nearly 60% of them expressed *Mchr1* mRNA (**Figure 4-4**) and were most likely found in the dorsomedial aspects of the VTA (**Figure 4D**). Taken together, these findings indicated the abundance of *Mchr1* mRNA throughout the VTA in glutamatergic, GABAergic, and/or dopaminergic VTA cells.

**Figure 4.**
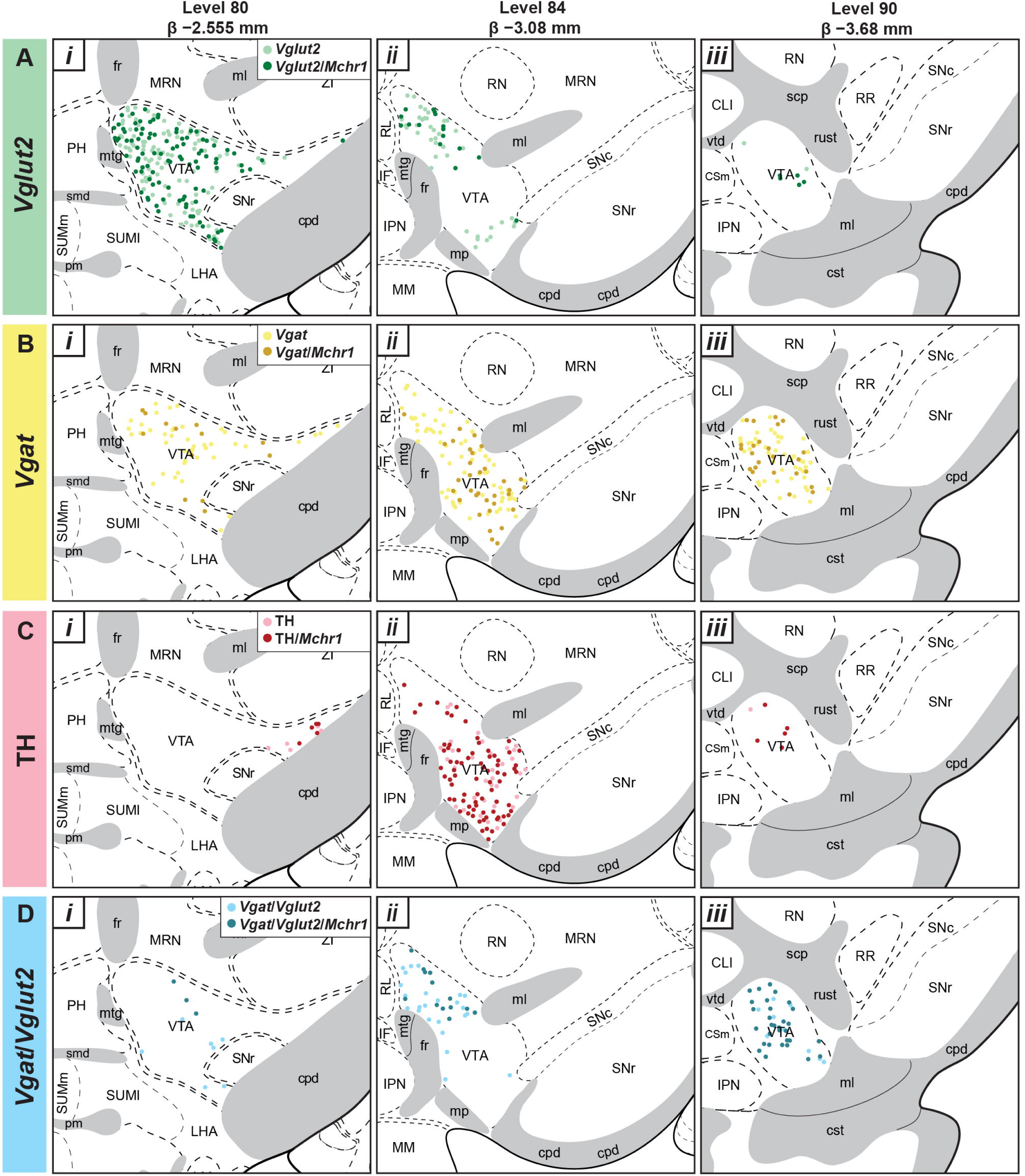
Spatial distribution of *Mchr1* mRNA in heterogeneous VTA cells. *Mchr1* mRNA hybridization expression in *Vglut2* (**A**; **Figure 4-1**), *Vgat* (**B**; **Figure 4-2**), TH (**C**; **Figure 4-3**), or dual *Vglut2* and *Vgat* VTA cells (**D**; Figure 4**-4**) mapped to a representative anterior (***i***), middle (***ii***), and posterior (***iii***) VTA section using *Allen Reference Atlas* (Dong, 2008) templates at the atlas level or Bregma value (β) indicated (*top*).

## MCH inhibited *Vgat* and *Th* VTA cells but not *Vglut2* VTA cells

Based on the spatial distribution of *Mchr1* expression in *Vglut2*, *Vgat*, and TH VTA cells (**Figure 4**), we performed whole-cell patch-clamp recordings in EGFP-labeled *Vglut2*^EGFP^, *Vgat*^EGFP^, and *Th*^EGFP^ VTA cells from *Vglut2-cre;L10-Egfp* (**Figure 5A*i***), *Vgat-cre;L10-Egfp* (**Figure 5B*i***), and *Th-cre;L10-Egfp* reporter mice (**Figure 5C*i***), respectively. MCH (3 μM) bath application had no effect on the resting membrane potential (baseline: −59.6 ± 3.2 mV; MCH: −59.7 ± 3.8 mV, n = 6 cells, t(5) = 0.20, p = 0.849, η^2^ = 0.01, paired t test) of *Vglut2*^EGFP^ VTA cells (n = 0/6 cells, N = 3; **Figure 5A*ii***). By contrast, MCH hyperpolarized *Vgat*^EGFP^ (baseline: −47.5 ± 2.3 mV; MCH: −52.9 ± 2.7 mV, n = 5/14 cells, N = 5, t(4) = 2.80, p = 0.049, η^2^ = 0.66, paired t test; **Figure 5B*ii***) and *Th*^EGFP^ cells (baseline: −53.5 ± 3.6 mV; MCH: −57.8 ± 2.8 mV, n = 6/16 cells, N = 12, t(5) = 3.15, p = 0.025, η^2^ = 0.66, paired t test; **Figure 5C*ii***) within 10 min and persisted for at least 25 min before gradually returning to baseline.

**Figure 5.**
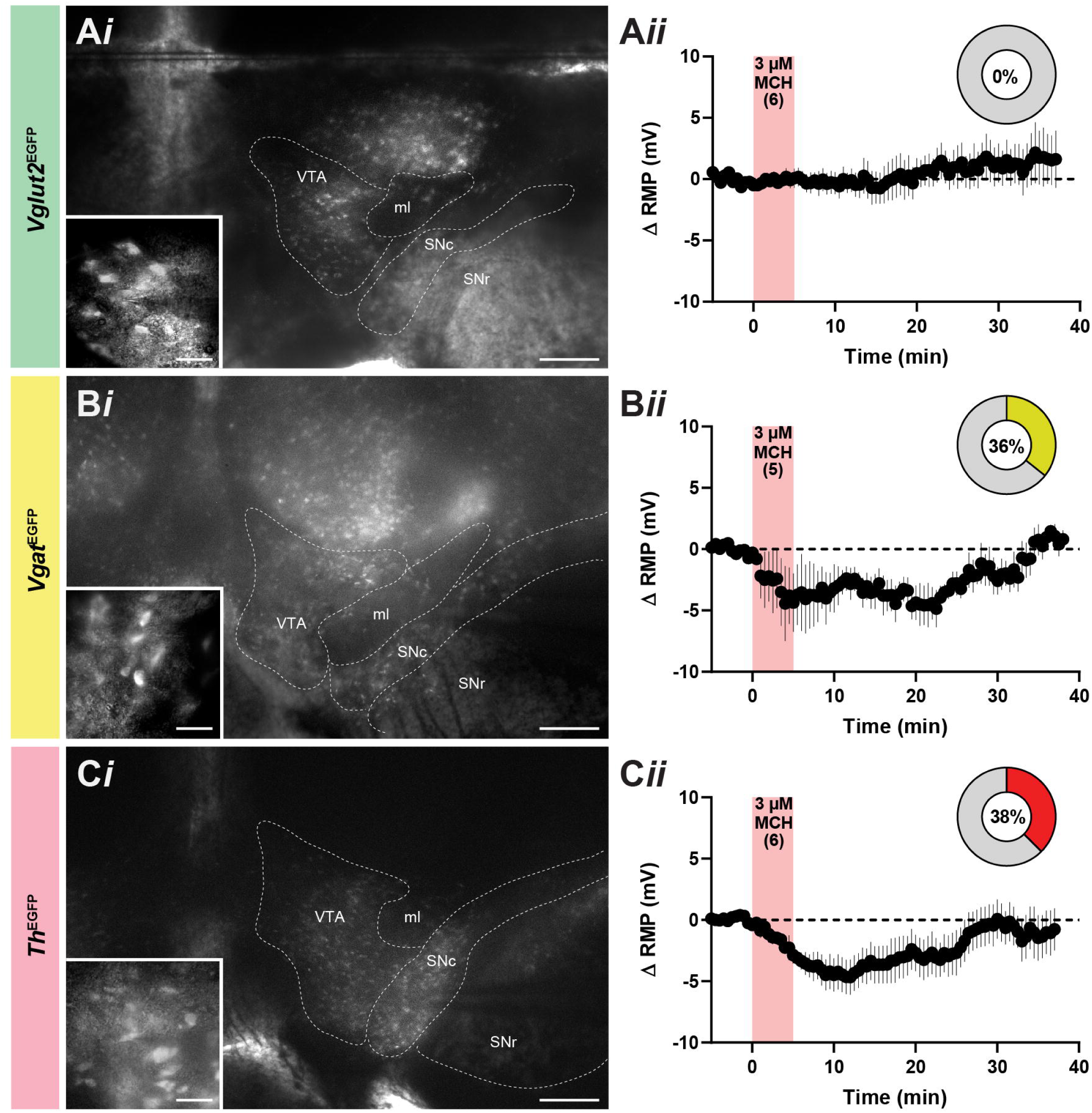
MCH hyperpolarized GABAergic and dopaminergic VTA cells. Representative epifluorescence image of coronal VTA brain slices (***i***) where cells expressing native EGFP fluorescence (inset in ***i***) from *Vglut2-cre;L10-Egfp* (**A**), *Vgat-cre;L10-Egfp* (**B**), and *Th-cre;L10-Egfp* reporter mice (**C**) guided whole-cell patch-clamp recordings to determine the change in resting membrane potential (Δ RMP) over time following the bath application of 3 μM MCH (***ii***) and proportion of MCH-responsive cells (donut plot in ***ii***). Scale bars: ***i***, 250 μm; *inset* in ***i***, 20 μm. ml, medial lemniscus; SNc, substantia nigra, compact part; SNr, substantia nigra, reticular part; VTA, ventral tegmental area.

While the proportion of MCH-sensitive *Vgat*^EGFP^ VTA cells could be predicted by the availability of *Mchr1* expression at *Vgat* cells (**Figure 4-2**), the proportion of MCH-sensitive *Th*^EGFP^ cells was lower than predicted by the prevalence of *Mchr1* expression at TH-ir cells (**Figure 4-3**). We first confirmed that over 80% (n = 13/16 cells, N = 7) of *Th*^EGFP^ cells in patch-clamp recordings expressed TH immunoreactivity (**Figure 6A**). We then determined the proportion of TH-ir cells that expressed MCHR1 protein (**Figure 6B**). Interestingly, only about one-third of TH-ir cells within the span of the VTA targeted for patch-clamp recordings (L84– 87) expressed MCHR1 (**Figure 6C**), which is comparable with a lower (38%) proportion of MCH-sensitive *Th*^EGFP^ cells surveyed by patch-clamp recordings (**Figure 5C*ii***).

**Figure 6.**
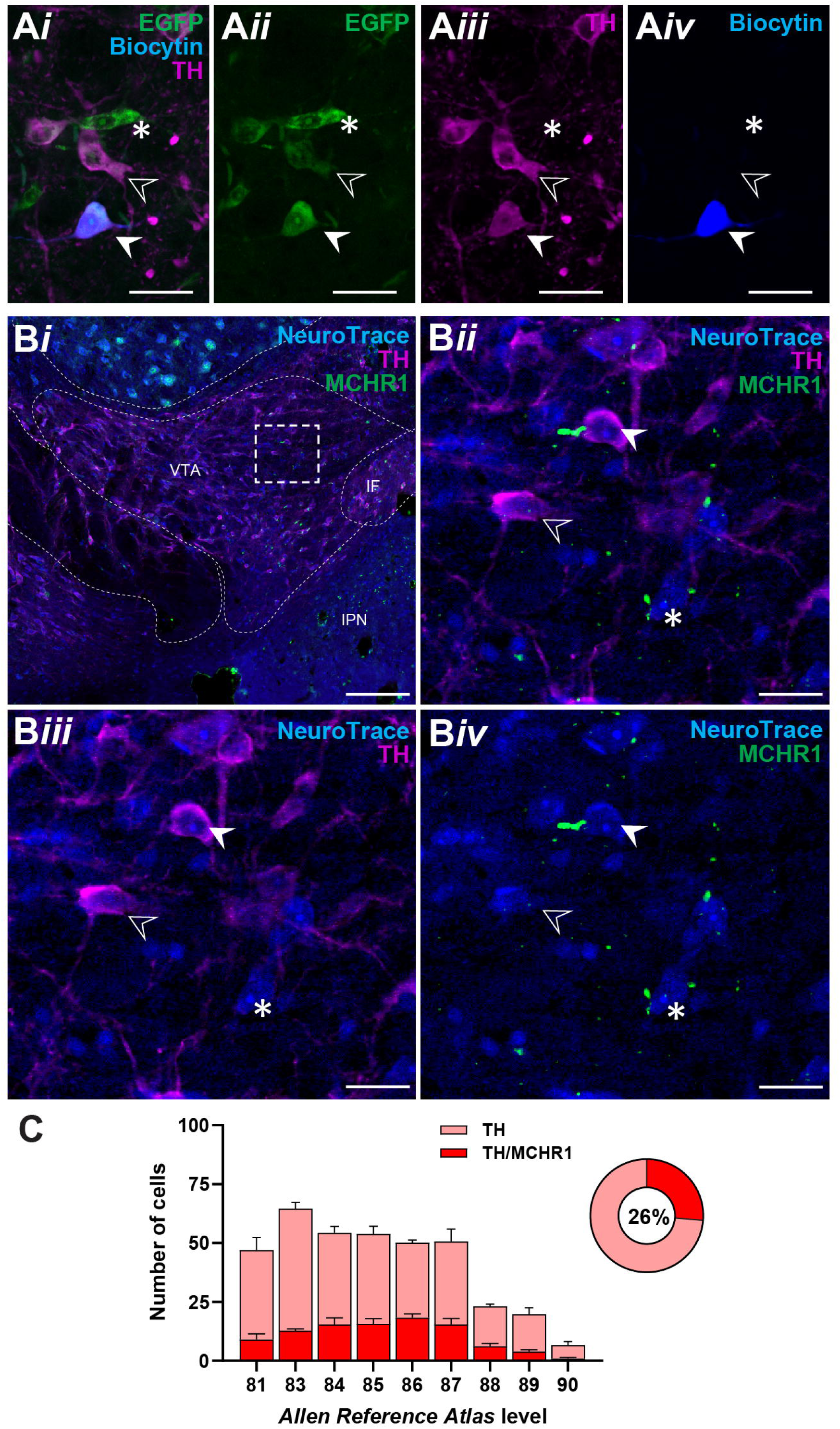
Selectivity of TH and MCHR1 coexpression in *Th*^EGFP^ VTA cells. Merged confocal photomicrograph (***i***) of native EGFP fluorescence (***ii***) and TH immunoreactivity (***iii***) in biocytin-filled (***iv***) cells from a subset of patch-clamp recordings from the VTA of *Th-cre;L10-Egfp* reporter mice (**A**). Merged confocal photomicrograph at low (***i***) and high magnification (***ii***, from dash-outlined region in ***i***) of TH- (***iii***) and MCHR1-immunoreactive signals (***iv***) on Nissl-stained (NeuroTrace) soma in wildtype VTA (**B**). Representative examples of the presence (white arrowhead) or absence (open arrowhead) of MCHR1 immunoreactivity at the primary cilium of VTA TH cells or non-TH-labeled cells (*). Number and proportion (donut plot) of MCHR1-positive (red) or -negative (pink) TH cells at each VTA level in accordance with the *Allen Reference Atlas* (*ARA*; Dong, 2008) of male and female wildtype mice (N = 4; **C**). Scale bars: **A**, 30 μm; **B*i***, 200 μm; **B*ii****–**iv***, 25 μm. IF, interfascicular nucleus raphe; IPN, interpeduncular nucleus; VTA, ventral tegmental area.

We then assessed if MCH-mediated inhibition of *Vgat*^EGFP^ and *Th*^EGFP^ VTA cells was activity-dependent by pretreatment with the voltage-gated sodium channel blocker tetrodotoxin (TTX). Since the effect of bath-applied MCH was so long-lasting, we first established if MCH-sensitive VTA cells could be identified by a short puff application of MCH (**Figure 7A*i***) lasting <5 seconds to allow rapid recovery of the resting membrane potential (**Figure 7A*ii***). A single puff of MCH produced an acute membrane hyperpolarization (baseline: −55.5 ± 4.2 mV; Δ RMP: −6.7 ± 1.8 mV, n = 4) that peaked within 10 seconds and returned to baseline within one minute. A second puff of MCH applied 3–5 minutes later at the same cell elicited a similar hyperpolarization (baseline: −55.9 ± 4.3 mV; Δ RMP: −4.3 ± 1.7 mV, n = 4) and time course (F(1, 6) = 3.3, p = 0.120, ANOVA; **Figure 7A*iii***). Importantly, puff application of ACSF produced visibly different outcomes than the first MCH puff (F(1, 5) = 6.8, p = 0.047, ANOVA) or second MCH puff (F(1, 5) = 30.7, p = 0.003, ANOVA), as it did not effect changes in membrane potential (baseline: −56.3 ± 2.7 mV; Δ RMP: −0.2 ± 0.6 mV, n = 3; **Figure 7A*iii***). This thus showed that a localized MCH puff could identify MCH-sensitive VTA cells by enabling short-lived but reproducible membrane hyperpolarizations that were not significantly desensitized by repeated application. We then puff-applied MCH to identify MCH-sensitive *Vgat*^EGFP^ and *Th*^EGFP^ VTA cells and compared the effect of MCH before and after TTX (500 nM) treatment. MCH-mediated hyperpolarization at *Vgat*^EGFP^ VTA cells (baseline: −56.3 ± 1.7 mV; Δ RMP: −1.8 ± 1.2 mV, n = 4) persisted in the presence of TTX (baseline: −55.2 ± 1.3 mV; Δ RMP: −1.1 ± 0.4 mV, n = 4) over time (F(1, 6) = 0.002, p = 0.965, ANOVA; **Figure 7B**). Similarly, *Th*^EGFP^ VTA cells remained sensitive to the peak effect of MCH (baseline: −57.1 ± 3.6 mV; Δ RMP: −5.4 ± 1.6 mV, n = 5) in the presence of TTX (baseline: −50.1 ± 3.4 mV; Δ RMP: −4.3 ± 1.0 mV, n = 5) and over time (F(1, 8) = 0.3, p = 0.623, ANOVA; **Figure 7C**). These findings indicated that MCH directly hyperpolarized *Vgat*^EGFP^ and *Th*^EGFP^ VTA cells.

**Figure 7.**
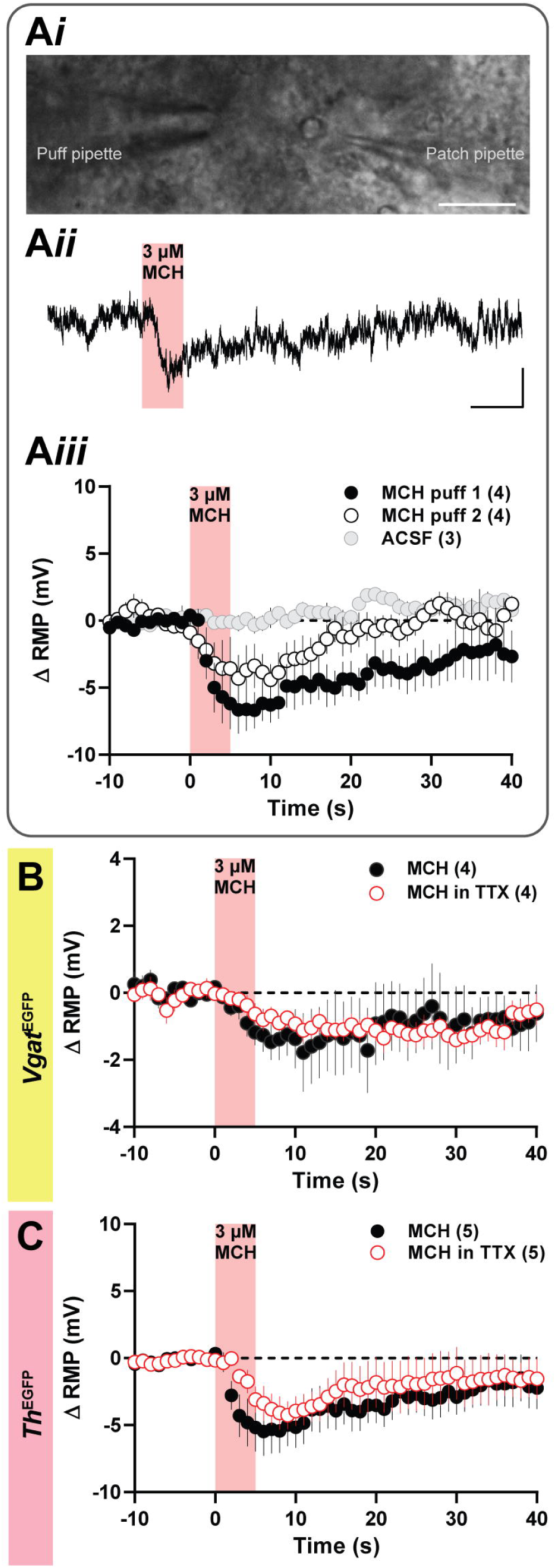
Rapid detection of direct MCH-mediated hyperpolarization at GABAergic and dopaminergic VTA cells. MCH-sensitive VTA cells identified by a localized puff of MCH applied within 20 μm of the patched cell (**A*i***). Representative sample trace of the membrane potential change following a puff of MCH (red shaded region) lasting <5 s (**A*ii***). Change in resting membrane potential (Δ RMP) over time following the first (black circles) and second localized puff (white circles) of 3 μM MCH or of bath ACSF (gray circles) onto a VTA cell (**A*iii***). RMP change over time following localized puff of 3 μM MCH in the presence (red outlined circles) or absence of 500 nM TTX (black circles) at *Vgat* ^EGFP^ (**B**) and *Th*^EGFP^ VTA cells (**C**). Scale bars: **A*i***, 25 μm; **A*ii***, 3 mV, 10 s.

## MCH increased glutamatergic input to *Th* VTA cells

Dopaminergic output from the VTA can be regulated by local glutamatergic and GABAergic circuitry (Bariselli et al., 2016; Morales & Margolis, 2017; Wang et al., 2015), thus we assessed if MCH regulated the GABAergic and/or glutamatergic input to *Th*^EGFP^ VTA cells. Bath application of MCH did not alter the frequency (baseline: 0.9 ± 0.1 Hz; MCH: 0.9 ± 0.1 Hz, n = 9, t(8) = 0.33, p = 0.750, η^2^ = 0.01, paired t test; **Figure 8A**, **B**) or amplitude (baseline: 13.1 ± 0.8 pA; MCH: 13.4 ± 0.8 pA, n = 9, t(8) = 0.60, p = 0.563, η^2^ = 0.04, paired t test; **Figure 8C**) of sIPSC events arriving at *Th*^EGFP^ VTA cells.

**Figure 8.**
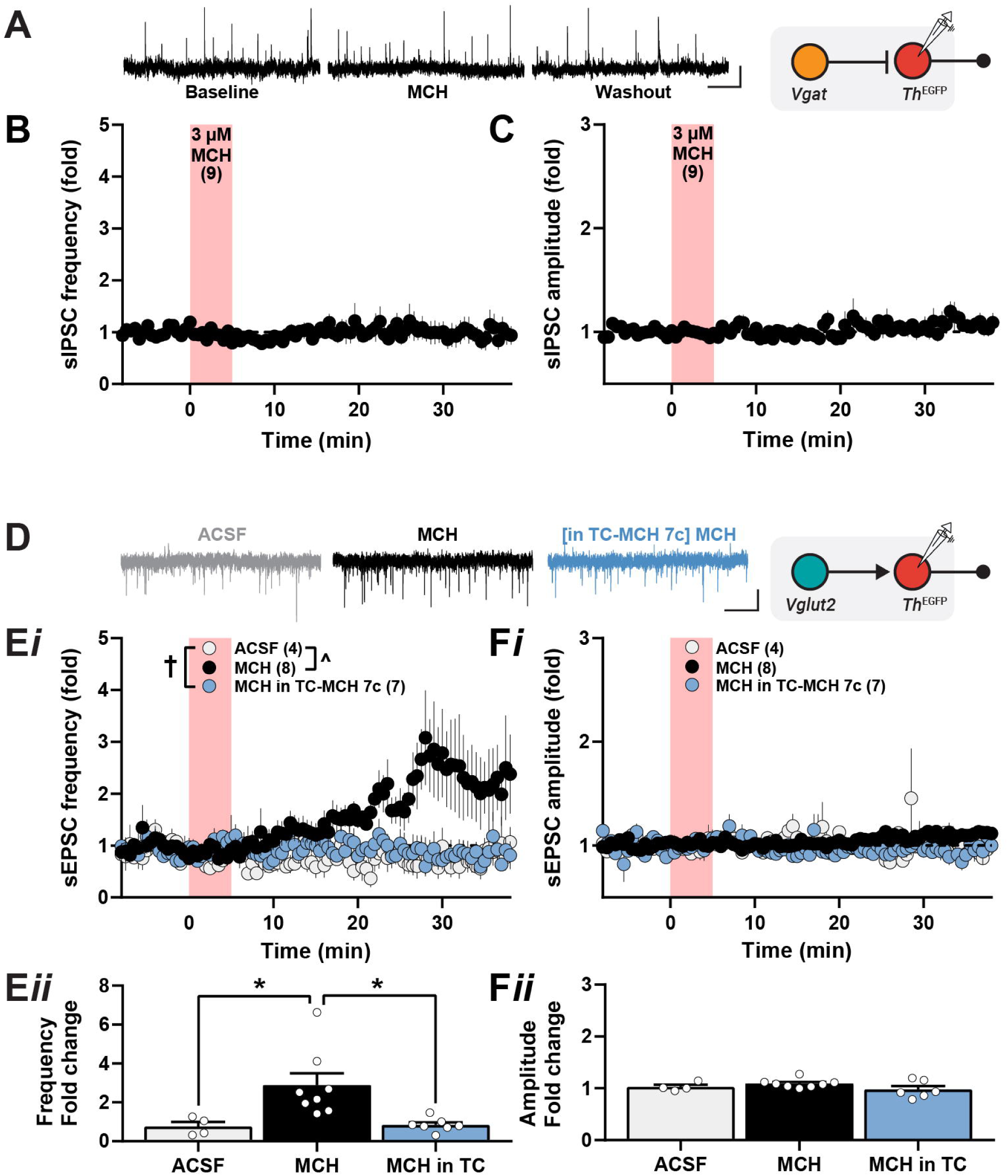
Delayed MCH-mediated increase of excitatory input to *Th*^EGFP^ VTA cells. Representative sample trace of spontaneous inhibitory postsynaptic current (sIPSC) events at baseline, following 3 μM MCH application, and MCH washout (*left*) from patch-clamp recordings at *Th*^EGFP^ cells (*right* schematic; **A**). Time course of the relative fold change in sIPSC frequency (**B**) and amplitude (**C**) relative to baseline values following bath application of 3 µM MCH. Representative sample trace of spontaneous excitatory postsynaptic current (sEPSC) events in ACSF, 3 μM MCH, and in MCH following pretreatment with 10 μM TC-MCH 7c (*left*) from patch-clamp recordings at *Th*^EGFP^ cells (*right* schematic; **D**). Time course of the relative (***i***) and peak (***ii***) fold change in sEPSC frequency (**E**) and amplitude (**F**) relative to baseline values following bath application of 3 µM MCH (black), bath ACSF only (gray), or MCH after pretreatment with 10 µM TC-MCH 7c (blue) for at least 10 minutes. One-way ANOVA: *, p < 0.05 or two-way ANOVA: †, p < 0.05 with Šidák multiple comparisons posttest: ^, p < 0.05. Scale bars: 10 pA, 2.5 s (**A**, **D**).

Interestingly, MCH elicited a delayed 3.1 ± 0.9-fold increase in baseline sEPSC frequency (1.3 ± 0.4 Hz, n = 8) at *Th*^EGFP^ VTA cells that gradually peaked about 25 minutes following MCH application (**Figure 8D**, **8E*i***). To determine if this delayed sEPSC frequency increase was MCH-dependent, we tracked the change in sEPSC frequency in the absence of MCH (ACSF only) or in the presence of a MCHR1 antagonist, TC-MCH 7c. Following MCH treatment, sEPSC frequency increased, from 1.3 ± 0.4 Hz to 2.6 ± 0.6 (n = 8, t(7) = 3.17, p = 0.016, η^2^ = 0.59, paired t test), which was not seen when bathed in ACSF only (baseline: 1.0 ± 0.4 Hz; MCH: 0.9 ± 0.5 Hz, n = 4, t(3) = 0.34, p = 0.760, η^2^ = 0.03, paired t test). The MCH-mediated increase in sEPSC frequency was also not seen when MCH was applied in the presence of TC-MCH 7c (baseline: 0.9 ± 0.4 Hz; MCH in TC-MCH 7c: 0.8 ± 0.3 Hz, n = 7, t(6) = 0.29, p = 0.780, η^2^ = 0.01, paired t test). There was a main effect of drug over time (F(2, 16) = 4.6, p = 0.027, ANOVA; **Figure 8E*i***), as the time course of fold change in sEPSC frequency with MCH treatment was significantly different than in ACSF (t(16) = 2.6, p = 0.036, Sidak post-test; **Figure 8E*i***) or when MCH was applied in the presence of TC-MCH 7c (t(15) = 2.4, p = 0.058, Sidak post-test; **Figure 8E*i***). This difference was largely attributed to a delayed increase in sEPSC frequency that was prominent >20 min after drug treatment. Overall, the fold change in sEPSC frequency in MCH (2.8 ± 0.6, n = 8) was significantly greater than in ACSF only (0.8 ± 0.2, n = 4; t(16) = 2.93, p = 0.029, ANOVA) or when applied at *Th*^EGFP^ VTA cells pretreated with TC-MCH 7c (0.8 ± 0.1, n = 7; t(16) = 3.33, p = 0.013, ANOVA; **Figure 8E*ii***). There was no main effect of MCH on sEPSC amplitudes over time (F(2, 16) = 2.3, p = 0.130; ANOVA; **Figure 8F*i***) or when comparing their fold change within each condition (F(2, 16) = 0.18, p = 0.176; one-way ANOVA; **Figure 8F*ii***).

To determine the role of upstream targets that facilitate excitatory input to *Th*^EGFP^ VTA cells, we isolated presynaptic excitatory input (mEPSC) with TTX pretreatment (**Figure 9A**). MCH-mediated glutamate release at *Th*^EGFP^ VTA cells was largely abolished in the presence of TTX (**Figure 9B**). However, a slight 1.5 ± 0.4-fold (baseline: 1.2 ± 0.5 Hz; MCH: 1.5 ± 0.6 Hz, n = 6; t(5) = 2.27, p = 0.072, η^2^ = 0.51, paired t test) increase in mEPSC frequency can be detected at some *Th*^EGFP^ VTA (**Figure 9B**), thus it is possible that MCH mediated glutamate release directly at glutamatergic terminals. There was no change in mEPSC amplitude following MCH treatment (baseline: 14.9 ± 1.9 pA; MCH: 14.9 ± 1.7 pA, n = 6; t(5) = 0.05, p = 0.961, η^2^ = 0.001, paired t test) or over time (**Figure 9C**). We thus considered that MCH may recruit upstream targets that indirectly regulate excitatory input at *Th*^EGFP^ VTA cells.

**Figure 9.**
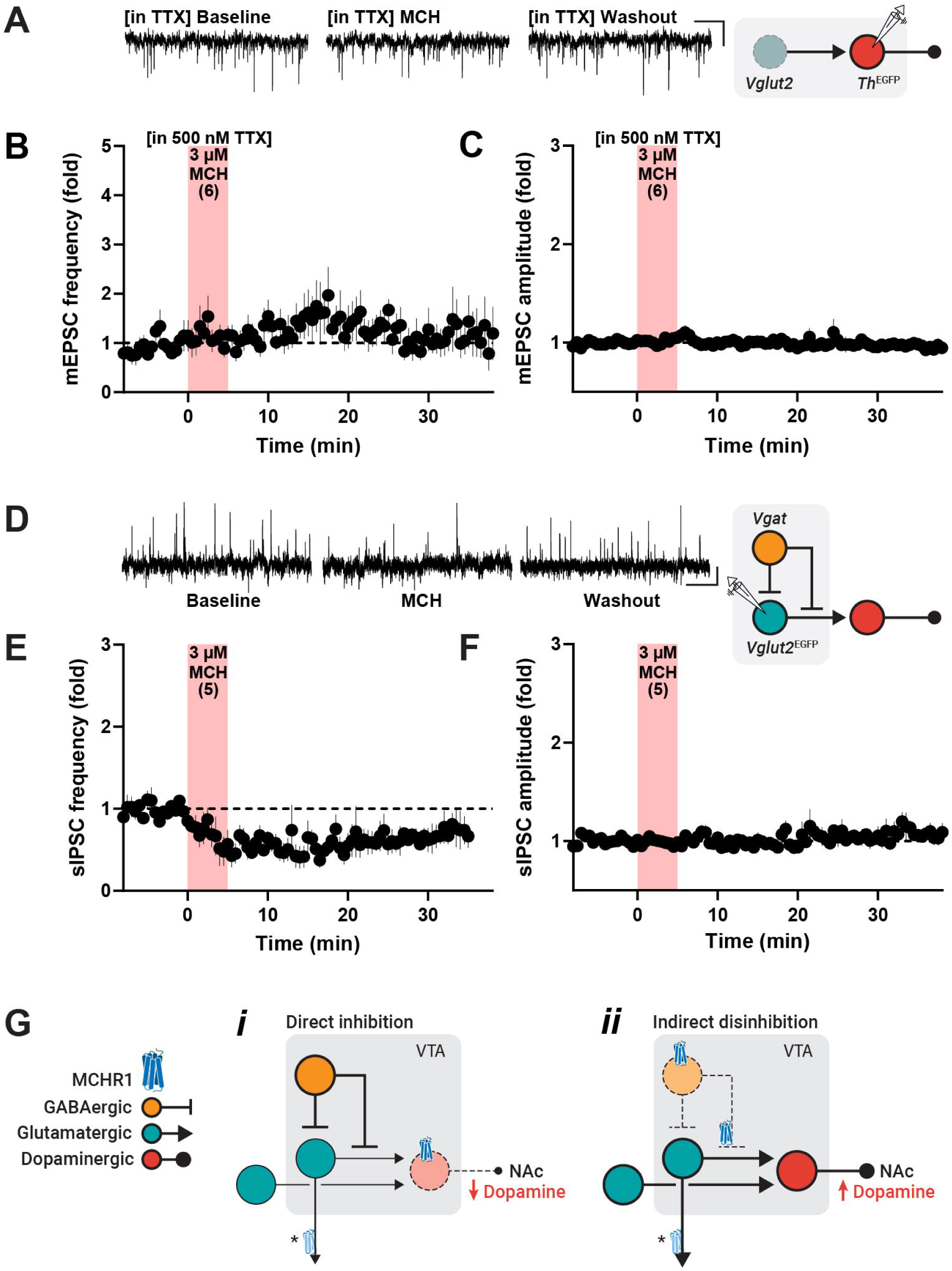
MCH-mediated disinhibition of local excitatory input to *Th* VTA cells. Representative sample trace of miniature excitatory postsynaptic current (mEPSC) events in the presence of 500 nM TTX at baseline, following 3 μM MCH application, and MCH washout (*left*) from patch-clamp recordings at *Th*^EGFP^ cells (*right* schematic; **A**). Time course of the relative fold change in mEPSC frequency (**B**) and amplitude (**C**) relative to baseline values following bath application of 3 µM MCH in slices pretreated with 500 nM TTX. Representative sample trace of spontaneous inhibitory postsynaptic current (sIPSC) events at baseline, following 3 μM MCH application, and MCH washout (*left*) from patch-clamp recordings at *Vglut2*^EGFP^ cells (*right* schematic; **D**). Time course of the relative fold change in sIPSC frequency (**E**) and amplitude (**F**) relative to baseline values following bath application of 3 µM MCH. Model of bidirectional, MCH-mediated regulation of dopamine release from the VTA to nucleus accumbens (NAc) via the mesolimbic pathway (**G**). Schematic showing inferred MCHR1 distribution within the VTA to elicit MCH-mediated direct inhibition of *Th*^EGFP^ VTA cells (***i***) and delayed, indirect GABA-mediated disinhibition of glutamatergic input to *Th*^EGFP^ VTA cells (***ii***). Putative MCHR1 expression (**\***) along efferent projections of VTA *Vglut2*^EGFP^ cells, which were not MCH-responsive. Scale bars: 5 pA, 2.5 s (**A**); 10 pA, 2.5 s (**D**).

As MCHR1 is a known inhibitory G_i/o_-protein coupled receptor (Hawes et al., 2000), we determined if the MCH-mediated increase in glutamatergic tone at *Th*^EGFP^ VTA cells may be ascribed to GABAergic disinhibition at glutamatergic nerve terminals (**Figure 9D**). We recorded from *Vglut2*^EGFP^ VTA cells and found that MCH produced a prolonged 0.4 ± 0.1-fold decrease in sIPSC frequency (baseline: 1.7 ± 0.4 Hz; MCH: 1.0 ± 0.3 Hz, n = 5; t(4) = 5.08, p = 0.007, η^2^ = 0.87, paired t test; **Figure 9E**) but not amplitude of sIPSC events (baseline: 14.4 ± 1.3 pA; MCH: 19.3 ± 5.0 pA, n = 5; t(4) = 0.95, p = 0.395, η^2^ = 0.19, paired t test; **Figure 9F**). Overall, these recordings suggested that, in addition to directly inhibiting *Th* VTA cells, MCH may also engage a local disynaptic circuit from *Vgat* and *Vglut2* VTA cells to regulate VTA output (**Figure 9G**).

## Discussion

MCH can directly and indirectly reglate *Th* VTA cells. MCHR1-expressing cells were widespread through the VTA, including at glutamatergic *Vglut2*, GABAergic *Vgat*, and dopaminergic TH cells, thus providing a neuroanatomical basis for MCH action in the VTA. Electrophysiological recordings showed that MCH hyperpolarized and inhibited *Th* cells and *Vgat* cells. Interestingly, MCH did not directly regulate *Vglut2* cells but disinhibited *Vglut2* afferents leading to increased excitatory transmission at *Th* cells. Therefore, functional MCHR1 activation in the VTA may provide bidirectional regulation of dopamine output from the VTA.

The majority, nearly 80%, of TH-ir VTA cells expressed *Mchr1* mRNA, and MCH directly hyperpolarized 38% of *Th* VTA cells in an activity-independent manner, which is consistent with MCHR1 protein expression at TH-ir VTA cells. Furthermore, it is possible that native EGFP fluorescence may not reflect active TH cells, as Cre expression in TH-negative VTA cells may comprise up to half of all EGFP*^Th^* cells (Lammel et al., 2015). However, when patching within the VTA, we found over 80% of patched EGFP*^Th^* cells expressed TH, indicating more robust EGFP*^Th^*expression in TH cells than previously reported.

MCHR1 signaling has known inhibitory actions (Chambers et al., 1999; Hawes et al., 2000; Lembo et al., 1999; Saito et al., 1999), including within the mesolimbic system connecting the VTA and ventral striatum. In the accumbens, MCH hyperpolarizes medium spiny neurons (Georgescu et al., 2005) and suppresses dopamine release (Chee et al., 2019). Furthermore, loss of MCH signaling increases accumbens dopamine accumulation leading to hyperactivity and increased energy expenditure (M. J. Chee et al., 2019; Mul et al., 2011; Pissios et al., 2008). As the main source of dopaminergic afferents to the accumbens arises from the VTA, it stands to reason that MCH may regulate dopamine release by direct inhibition of VTA neurons.

Interestingly, MCH action within the VTA caused a delayed increase in excitatory input to *Th* VTA cells. MCHR1 activation engages G_i/o_- or G_q_-protein systems (Pissios et al., 2003; Sears et al., 2010) to elicit inhibitory or excitatory actions, respectively. We considered that MCH may act presynaptically to stimulate glutamatergic afferents, but the MCH-mediated increase in glutamatergic transmission was abolished in the presence of TTX. This suggested that the stimulatory effect of MCH on glutamatergic afferents was indirect, activity-dependent, and polysynaptic. We thus examined if MCH may indirectly recruit glutamatergic afferents detected at *Th* VTA cells by suppressing GABAergic input onto *Vglut2* VTA cells or axons. We found that MCH directly inhibited GABAergic VTA cells and suppressed GABAergic input to *Vglut2* VTA cells. In effect, MCH can suppress GABAergic activity to disinhibit glutamatergic transmission at *Th* VTA cells, and we posit that this GABA-dependent microcircuit within the VTA may serve to restore the output of *Th* VTA cells that were acutely inhibited by MCH (**Figure 9G**).

VTA *Vglut2* cells are known to innervate (Dobi et al., 2010) and regulate dopaminergic VTA cells (Bariselli et al., 2016; Wang et al., 2015). MCH had no effect on the membrane potential of *Vglut2* VTA cells. *Vglut2* cells were more abundant in the anterior VTA (Morales & Root, 2014; Root et al., 2018; Yamaguchi et al., 2007), thus it is possible that the null effect of MCH at *Vglut2* cells is ascribed to differences in electrophysiological effects in the anterior and posterior VTA (Guan et al., 2012). It may thus be insightful to track the rostrocaudal distribution of subsequent patch-clamp recordings. By contrast, MCH increased glutamatergic tone at *Th* VTA cells. *Vglut2* VTA cells form local synapses with dopaminergic and non-dopaminergic VTA neurons (Dobi et al., 2010) and are pivotal regulators of local VTA circuitry and output. Glutamatergic VTA cells can drive dopamine release and promote conditioned place preference (Wang et al., 2015) or drug-induced conditioned place preference (Mead & Stephens, 1999; Slusher et al., 2001; Tzschentke & Schmidt, 1998) that can be enhanced by food restriction (Stuber et al., 2002) to assess drug reward (Hiroi & White, 1991). Interestingly, *Mchr1* deletion can block the effect of food restriction on amphetamine-induced conditioned place preference (Geuzaine et al., 2014). However, as *Vglut2* cells within the VTA were not MCH-sensitive, it would be important to consider that MCH may indirectly modulate dopamine output at glutamatergic afferents arising from outside the VTA, such as from the prefrontal cortex (Beier et al., 2015) or lateral hypothalamus (Geisler & Wise, 2008; Watabe-Uchida et al., 2012). Furthermore, since we did not detect postsynaptic MCH effects at the soma of *Vglut2* cells, we predicted that MCHR1 distribution within *Vglut2* cells would accumulate at presynaptic terminals to regulate glutamate release at projection sites (**Figure 9G**, *asterisk*).

Prominent *Vgat* expression in the posterior VTA may partly form the tail of the VTA (tVTA) or rostromedial tegmental nucleus (RMTg; Jhou et al., 2009). The GABAergic cell cluster in the posterior VTA also reflected the highest proportion of *Vgat* VTA cells that expressed *Mchr1*. This supports the hypothesis that the posterior VTA is anatomically and functionally unique (Lahti et al., 2016; Smith et al., 2019) and suggests that this subregion is a site of MCH action. Sex may interact with *Vgat* expression at *Mchr1* cells in the VTA, as there is a larger portion of *Mchr1* cells expressing *Vgat* in the VTA of female mice. Therefore, MCH may contribute to sex-specific regulation of VTA output, potentially via *Vgat* cells in females or by yet characterized *Mchr1*-only cells in males. The tVTA/RMTg has been implicated in responsiveness to alcohol as exposure to alcohol stimulates tVTA/RMTg neurons, and rats will self-administer ethanol to the posterior but not anterior VTA (Guan et al., 2012). Interestingly, *Mchr1* deletion suppresses alcohol-induced conditioned place preference (Karlsson et al., 2016), and MCHR1 blockade decreases self-administration of alcohol (Cippitelli et al., 2010). The posterior VTA supplies dense GABAergic projections to other VTA neurons (Barrot et al., 2012; Jhou et al., 2009; Lecca et al., 2012) and provides strong inhibitory tone to dopaminergic VTA neurons (Barrot et al., 2012; Bouarab et al., 2019; Guan et al., 2012). While MCH did not regulate GABAergic synaptic transmission at *Th* VTA cells, it inhibited GABAergic transmission at glutamatergic cells. Furthermore, MCH directly inhibited *Vgat* cells, thus, taken together, MCH action at *Vgat* cells in the posterior VTA may weaken inhibition of *Th* VTA cells to indirectly facilitate dopaminergic output.

Consistent with findings in rats (Hervieu et al., 2000; Saito et al., 2001), the relative expression of *Mchr1* mRNA in the VTA was comparable to that in the hypothalamus and hippocampus. *Mchr1* expression was visible in all major subtypes of VTA cells, including cells that coexpressed two or more chemical messengers. *Vgat*/*Vglut2* cells were the most common subpopulation coexpressing a combination of TH, *Vgat*, or *Vglut2* and is consistent with previous reports of GABA and glutamate corelease from VTA cells (Root et al., 2018; Root, Mejias-Aponte, Zhang, et al., 2014). VTA neurons may also be both glutamatergic and dopaminergic (Morales & Root, 2014; Morales & Margolis, 2017; Yamaguchi et al., 2011), but unlike the rat VTA, the incidence of colocalization between TH and *Vglut2* was low. In rats, most dopaminergic and glutamatergic VTA cells were at or very close to the midline (Yamaguchi et al., 2011) and includes midline regions encompassing the interfascicular nucleus raphe, rostral linear nucleus raphe, and central linear nucleus raphe as subregions of the VTA (Morales & Root, 2014; Morales & Margolis, 2017; Yamaguchi et al., 2011). However, our analyses were based on the neuroanatomical boundaries of the VTA as defined by the *Allen Reference Atlas* (Dong, 2008) and so did not include midline outside the VTA where dual TH/*Vglut2* cells may be most abundant. Finally, although VTA cells may co-release dopamine and GABA (Stamatakis et al., 2013; Tritsch et al., 2012), coexpression of TH and *Vgat* did not form a substantial population within the VTA. GABA release does not require vesicular transport via vGAT because the vesicular monoamine transporter 2 (vMAT2) can fully support GABA release from dopaminergic cells (Tritsch et al., 2012), thus it is not surprising that TH/*Vgat* coexpression was low.

In conclusion, our findings suggest that MCHR1 expression at multiple VTA cell types can modulate the local VTA microcircuit to provide bidirectional regulation of dopamine output from the VTA. MCH directly inhibited dopaminergic VTA cells, which may acutely suppress dopamine release. However, MCH also inhibited GABAergic VTA cells, which may disinhibit glutamatergic input to dopamine cells. This GABA-mediated disinhibition has a delayed time course and may act to restore dopamine levels acutely lowered by the direct MCH-mediated inhibition of dopamine cells. Paradoxically, this may be in contrast to prior findings that *Mchr1* deletion from *Vgat* neurons results in a hyperdopaminergic state (Chee et al., 2019), but this and previous findings may not be mutually exclusive, as MCH can have site-specific effects at different nodes of the mesolimbic dopamine pathway. It would be of interest to test our proposed model (**Figure 9G**) in future studies and determine if *Mchr1* deletion from dopaminergic cells would also elicit a hyperdopaminergic state, or whether disrupting GABAergic transmission would abolish the facilitation of glutamatergic input to dopamine cells.

## Conflicts of Interest

The authors have no conflicts of interest to declare.

## Acknowledgements

This study is supported by Natural Sciences and Engineering Research Council of Canada (NSERC) Discovery Grant RGPIN-2017-06272 (MJC), NSERC Canadian Graduate Master’s Scholarship and the Ontario QEII-GSST program (CDS), Carleton University’s Internship-Carleton University Research Experience for Undergraduate Students (I-CUREUS) (PAM, JWI).

## Extended Data

**Figure 3-1. Relative distribution of *Mchr1*-expressing VTA cells** in male and female mice (**A**). Sex may interact with *Vgat* expression to shift the distribution of *Mchr1* mRNA (**B**).

**Figure 4-1. Distribution of *Mchr1*-expressing *Vglut2* VTA cells.** Number of *Mchr1*-positive (dark green) or -negative (light green) *Vglut2* cells relative to all VTA cells counted (gray) at each level in accordance with the *Allen Reference Atlas* (*ARA*; Dong, 2008) (N = 6; **A**). Presence (dark green circles) or absence (light green circles) of *Mchr1* mRNA hybridization in *Vglut2* cells throughout the anteroposterior extent of the VTA (**B**–**L**) mapped to *ARA*. Each panel includes the nomenclature, atlas level and Bregma value (β) assigned to that tissue in accordance with the *ARA*.

**Figure 4-2. Distribution of *Mchr1*-expressing *Vgat* VTA cells.** Number of *Mchr1*-positive (dark yellow) or -negative (light yellow) *Vgat* cells relative to all VTA cells counted (gray) at each level in accordance with the *Allen Reference Atlas* (*ARA*; Dong, 2008) (N = 6; **A**). Presence (dark yellow circles) or absence (light yellow circles) of *Mchr1* mRNA hybridization in *Vgat* cells throughout the anteroposterior extent of the VTA (**B**–**L**) mapped to *ARA*. Each panel includes the nomenclature, atlas level and Bregma value (β) assigned to that tissue in accordance with the *ARA*.

**Figure 4-3. Distribution of *Mchr1*-expressing TH VTA cells.** Number of *Mchr1*-positive (red) or -negative (pink) TH-immunoreactive cells relative to all VTA cells counted (gray) at each level in accordance with the *Allen Reference Atlas* (*ARA*; Dong, 2008) (N = 6; **A**). Presence (red circles) or absence (pink circles) of *Mchr1* mRNA hybridization in TH cells throughout the anteroposterior extent of the VTA (**B**–**L**) mapped to *ARA*. Each panel includes the nomenclature, atlas level and Bregma value (β) assigned to that tissue in accordance with the *ARA*.

**Figure 4-4. Distribution of *Mchr1* mRNA expression *Vgat*-and *Vglut2*-positive VTA cells.** Number of *Mchr1*-positive (dark blue) or -negative (light blue) cells expressing both *Vglut2* and *Vgat* mRNA relative to all VTA cells counted (gray) at each level in accordance with the *Allen Reference Atlas* (*ARA*; Dong, 2008) (N = 6; **A**). Presence (dark blue circles) or absence (light blue circles) of *Mchr1* mRNA hybridization in dual-labeled *Vglut2* and *Vgat* cells throughout the anteroposterior extent of the VTA (**B**–**L**) mapped to *ARA*. Each panel includes the nomenclature, atlas level and Bregma value (β) assigned to that tissue in accordance with the *ARA*.

## Notes

### Competing Interest Statement

The authors have declared no competing interest.

### Summary of Updates

In this revision, we added new experiments or analyses focused on elucidating the disinhibition of dopamine cells by showing: -Experimental details on VTA slices and availability of all cell types in our slices -Inclusion of effect size (eta-squared) indicators, as appropriate -No change in the amplitude of synaptic events thus ruling out effects of postsynaptic origin -Lack of direct MCH action at glutamatergic afferents (via TTX pretreatment) -MCHR1-dependent increase in glutamate tone at dopaminergic cells (via MCHR1 antagonism) -MCH suppressed GABA tone at local glutamatergic cells in VTA -Schematic model integrating both acute inhibition and delayed disinhibition at dopaminergic cells

