## Supplementary figures and images for "Regulation of the mouse ventral tegmental area by melanin-concentrating hormone"

### Supplemental Figure 3-1

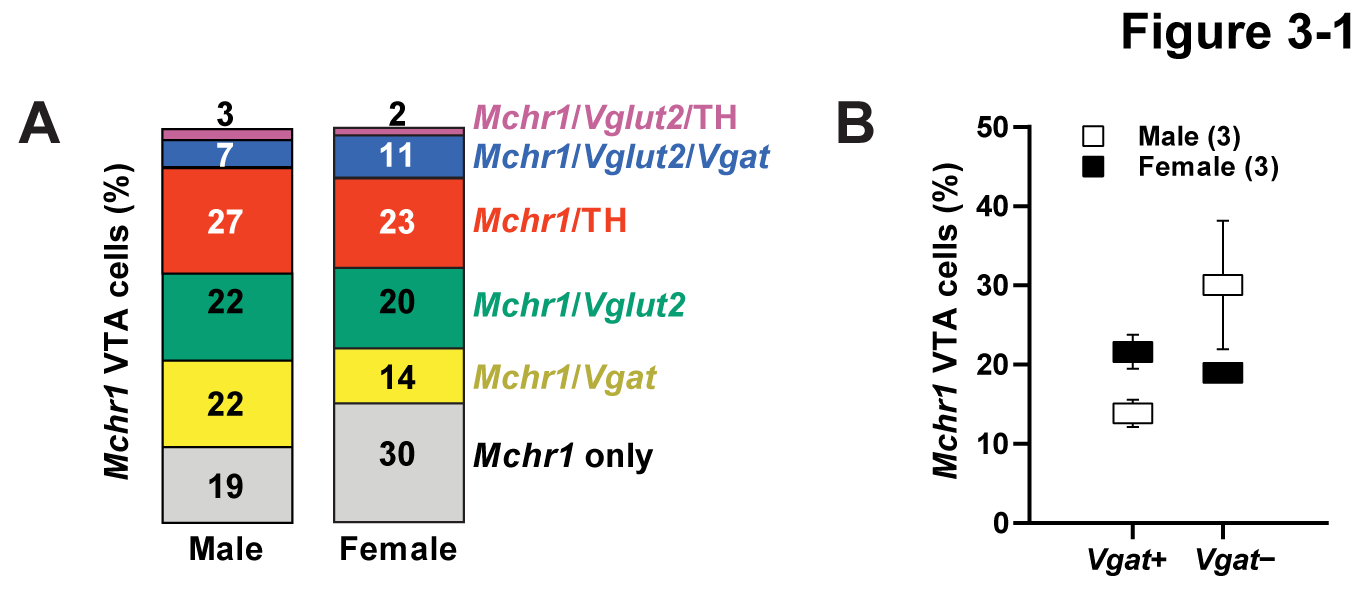

### Supplemental Figure 4-1

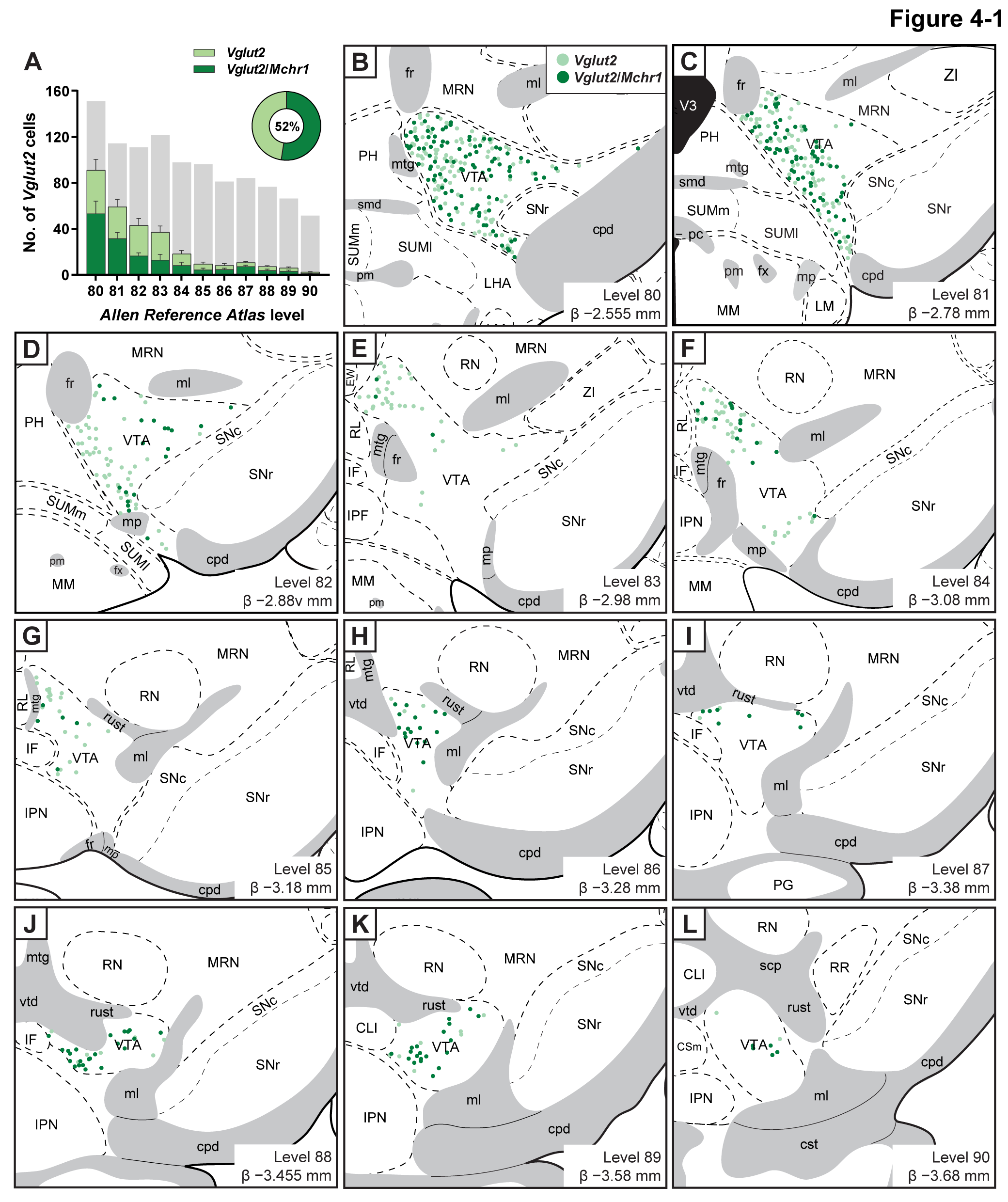

### Supplemental Figure 4-2

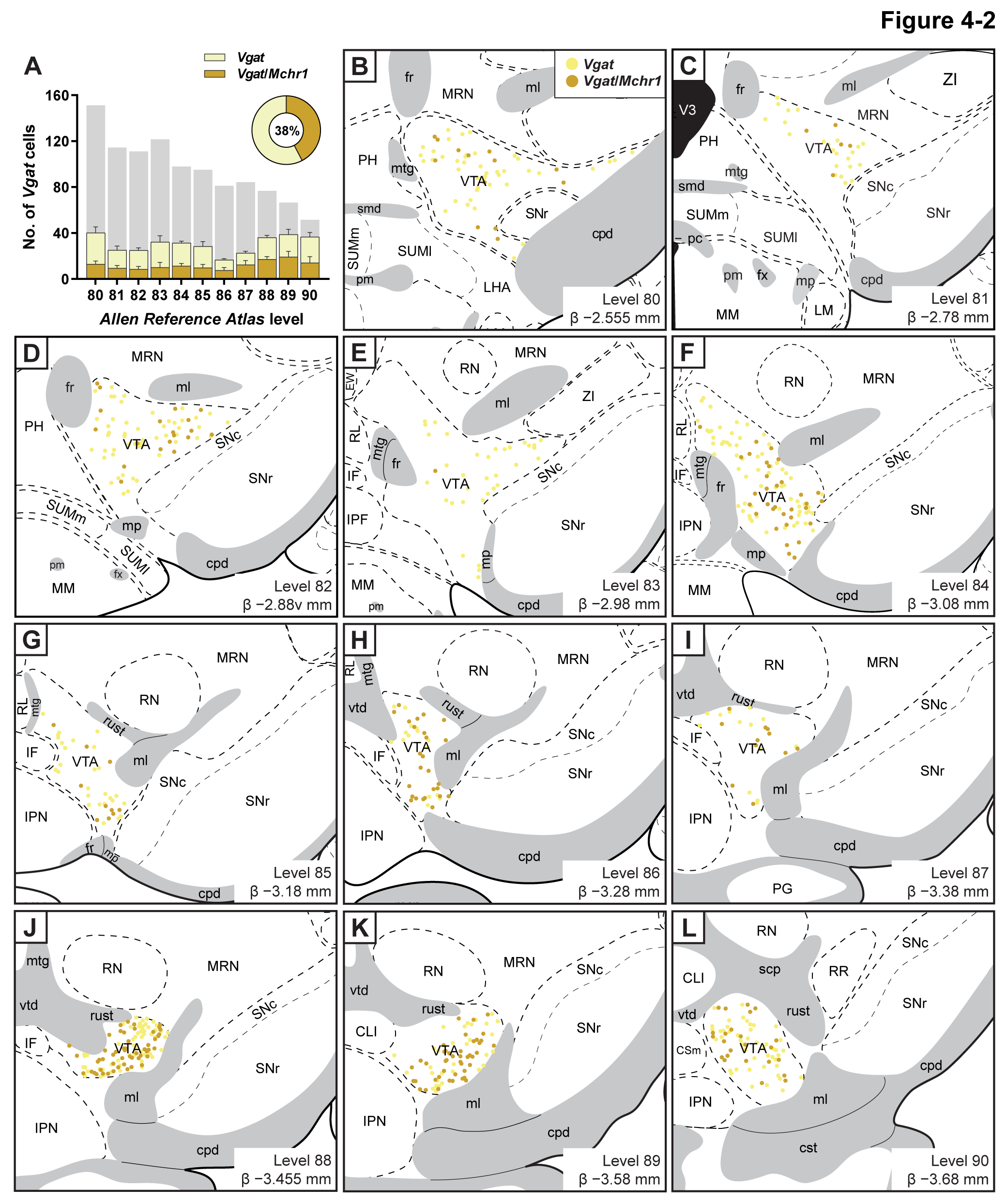

### Supplemental Figure 4-3

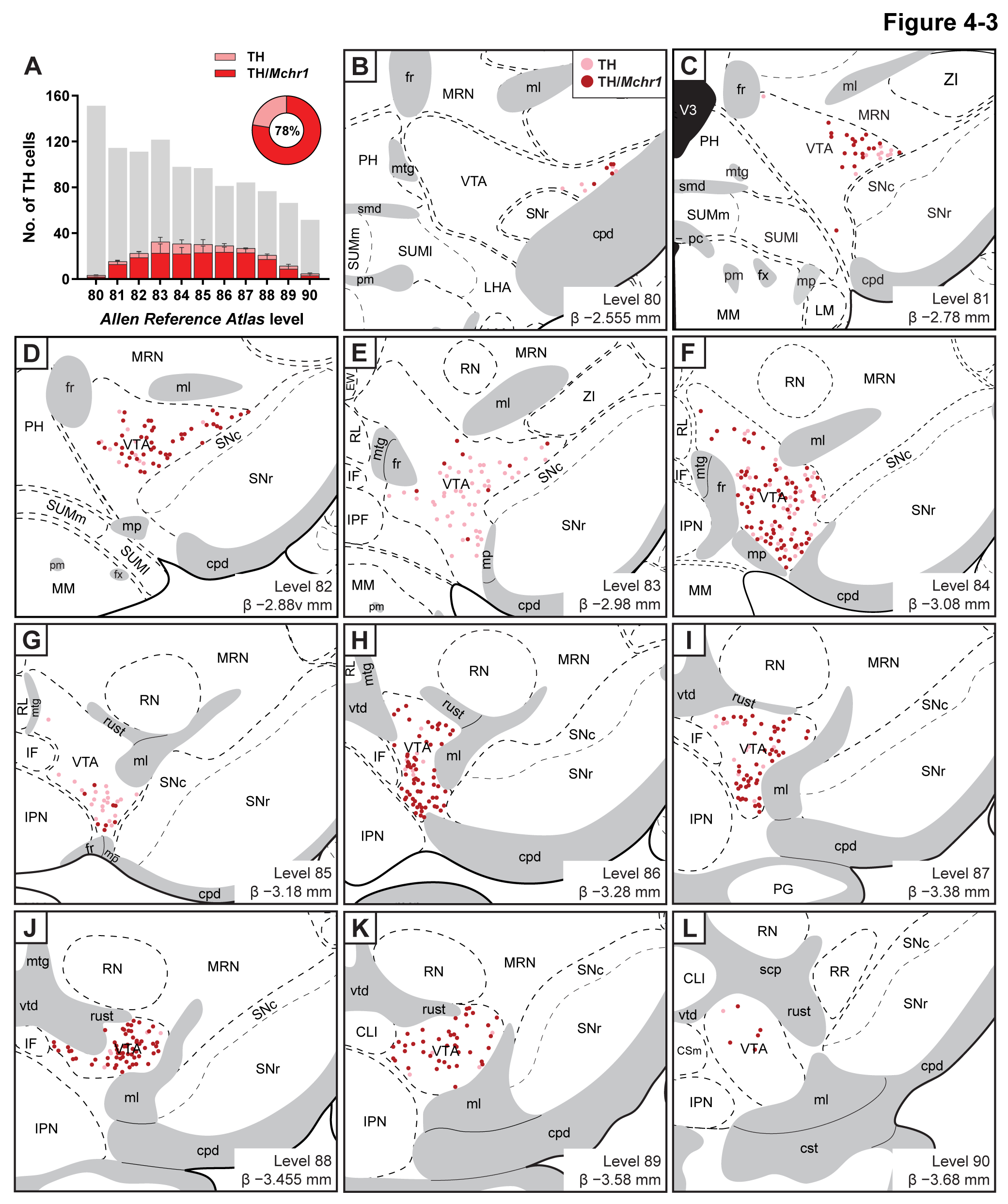

### Supplemental Figure 4-4

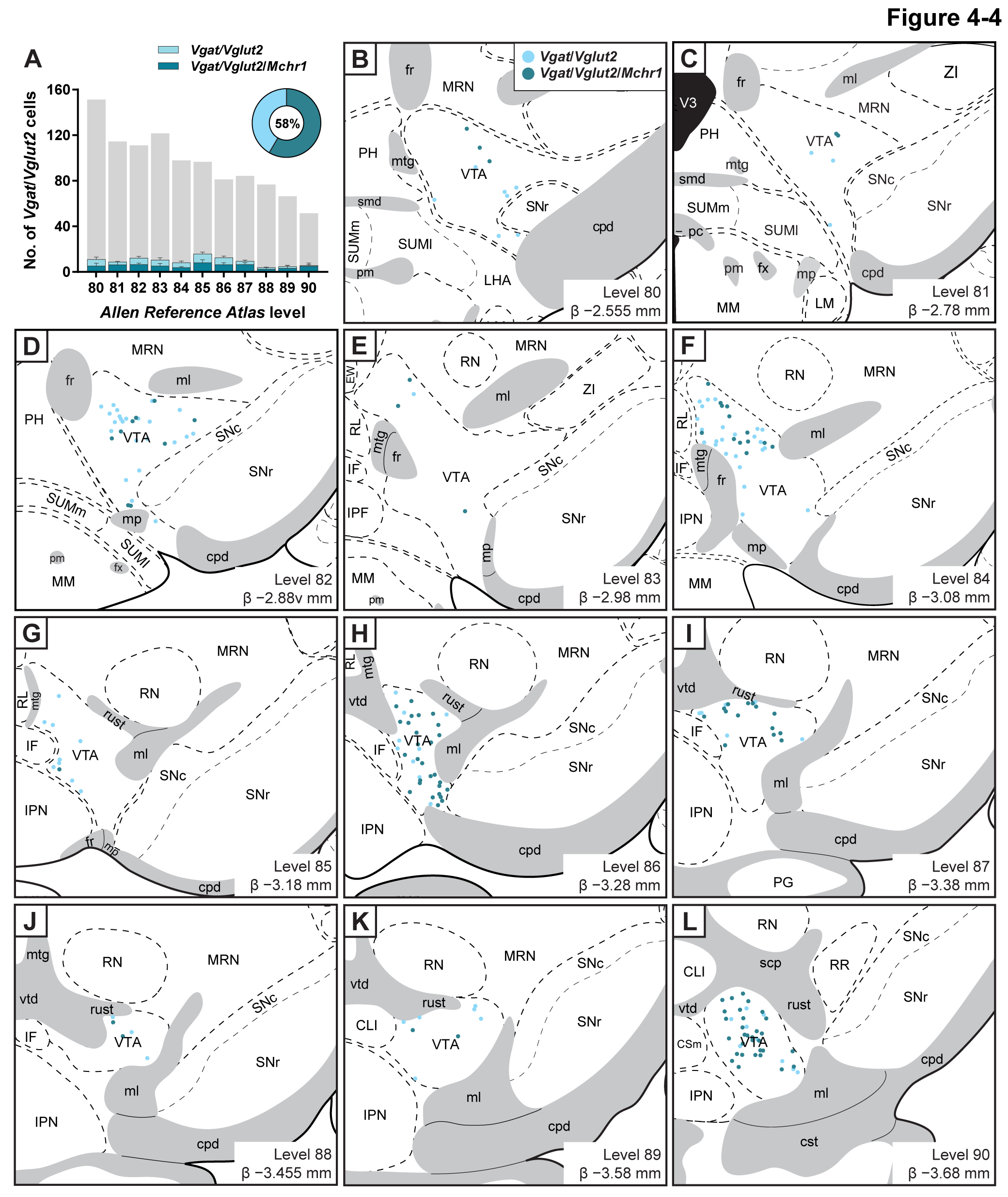
